# Exome sequencing identifies high-impact trait-associated alleles enriched in Finns

**DOI:** 10.1101/464255

**Authors:** Adam E Locke, Karyn Meltz Steinberg, Charleston WK Chiang, Susan K Service, Aki S Havulinna, Laurel Stell, Matti Pirinen, Haley J Abel, Colby C Chiang, Robert S Fulton, Anne U Jackson, Chul Joo Kang, Krishna L Kanchi, Daniel C Koboldt, David E Larson, Joanne Nelson, Thomas J Nicholas, Arto Pietilä, Vasily Ramensky, Debashree Ray, Laura J Scott, Heather M Stringham, Jagadish Vangipurapu, Ryan Welch, Pranav Yajnik, Xianyong Yin, Johan G Eriksson, Mika Ala-Korpela, Marjo-Riitta Järvelin, Minna Männikkö, Hannele Laivuori, FinnGen Project, Susan K Dutcher, Nathan O Stitziel, Richard K Wilson, Ira M Hall, Chiara Sabatti, Aarno Palotie, Veikko Salomaa, Markku Laakso, Samuli Ripatti, Michael Boehnke, Nelson B Freimer

**Affiliations:** Department of Medicine, Washington University School of Medicine, St. Louis, MO; McDonnell Genome Institute, Washington University School of Medicine, St. Louis, MO; Department of Biostatistics and Center for Statistical Genetics, University of Michigan School of Public Health, Ann Arbor, MI; Department of Pediatrics, Washington University School of Medicine, St. Louis, MO; Center for Neurobehavioral Genetics, Jane and Terry Semel Institute for Neuroscience and Human Behavior, University of California Los Angeles, Los Angeles, CA; Center for Genetic Epidemiology, Department of Preventive Medicine, Keck School of Medicine, University of Southern California, Los Angeles, CA; Institute for Molecular Medicine Finland (FIMM), University of Helsinki, Helsinki, Finland; National Institute for Health and Welfare, Helsinki, Finland; Department of Biomedical Data Science, Stanford University, Stanford, CA; Department of Public Health, University of Helsinki, Helsinki, Finland; Helsinki Institute for Information Technology HIIT and Department of Mathematics and Statistics, University of Helsinki, Helsinki, Finland; Department of Genetics, Washington University School of Medicine, St. Louis, MO; The Institute for Genomic Medicine, Nationwide Children’s Hospital, Columbus, OH; Department of Pediatrics, The Ohio State University College of Medicine, Columbus, OH; USTAR Center for Genetic Discovery and Department of Human Genetics, University of Utah, Salt Lake City, UT; Federal State Institution “National Medical Research Center for Preventive Medicine” of the Ministry of Healthcare of the Russian Federation, Moscow, Russia; Departments of Epidemiology and Biostatistics, Bloomberg School of Public Health, Johns Hopkins University, Baltimore, MD; Institute of Clinical Medicine, Internal Medicine, University of Eastern Finland, Kuopio, Finland; Department of Public Health Solutions, National Institute for Health and Welfare, Helsinki, Finland; Folkhälsan Research Center, Helsinki, Finland; Department of General Practice and Primary Health Care, University of Helsinki, Helsinki and Helsinki University Hospital, Helsinki, Finland; Systems Epidemiology, Baker Heart and Diabetes Institute, Melbourne, Victoria, Australia; Computational Medicine, Faculty of Medicine, University of Oulu and Biocenter Oulu, University of Oulu, Oulu, Finland; NMR Metabolomics Laboratory, School of Pharmacy, University of Eastern Finland, Kuopio, Finland; Population Health Science, Bristol Medical School, University of Bristol, Bristol, UK; Medical Research Council Integrative Epidemiology Unit at the University of Bristol, Bristol, UK; Department of Epidemiology and Preventive Medicine, School of Public Health and Preventive Medicine, Faculty of Medicine, Nursing and Health Sciences, The Alfred Hospital, Monash University, Melbourne, Victoria, Australia; Biocenter Oulu, University of Oulu, Oulu, Finland; Center for Life Course Health Research, Faculty of Medicine, University of Oulu, Oulu, Finland; Unit of Primary Health Care, Oulu University Hospital, Oulu, Finland; Department of Epidemiology and Biostatistics, MRC-PHE Centre for Environment and Health, School of Public Health, Imperial College London, London, UK; Department of Life Sciences, College of Health and Life Sciences, Brunel University London, Uxbridge, UK; Northern Finland Birth Cohorts, Faculty of Medicine, University of Oulu, Oulu, Finland; Medical and Clinical Genetics, University of Helsinki and Helsinki University Hospital, Helsinki, Finland; Department of Obstetrics and Gynecology, Tampere University Hospital and University of Tampere, Faculty of Medicine and Life Sciences, Tampere, Finland; Cardiovascular Division, Department of Medicine, Washington University School of Medicine, St. Louis, MO; Department of Statistics, Stanford University, Stanford, CA; Analytical and Translational Genetics Unit (ATGU), Psychiatric & Neurodevelopmental Genetics Unit, Departments of Psychiatry and Neurology, Massachusetts General Hospital, Boston, MA; Broad Institute of MIT and Harvard, Cambridge, MA; Department of Medicine, Kuopio University Hospital, Kuopio, Finland

## Abstract

As yet undiscovered rare variants are hypothesized to substantially influence an individual’s risk for common diseases and traits, but sequencing studies aiming to identify such variants have generally been underpowered. In isolated populations that have expanded rapidly after a population bottleneck, deleterious alleles that passed through the bottleneck may be maintained at much higher frequencies than in other populations. In an exome sequencing study of nearly 20,000 cohort participants from northern and eastern Finnish populations that exemplify this phenomenon, most novel trait-associated deleterious variants are seen only in Finland or display frequencies more than 20 times higher than in other European populations. These enriched alleles underlie 34 novel associations with 21 disease-related quantitative traits and demonstrate a geographical clustering equivalent to that of Mendelian disease mutations characteristic of the Finnish population. Sequencing studies in populations without this unique history would require hundreds of thousands to millions of participants for comparable power for these variants.

## INTRODUCTION

Genotyping studies of common genetic variants (defined here as minor allele frequency [MAF]>1%) have identified tens of thousands of genome-wide significant associations with common diseases and disease-related quantitative traits^1^. For most traits, however, these associations account for only a modest fraction of trait heritability, and the mechanisms through which associated variants contribute to biological processes remain mostly unknown. These observations have led to the expectation that rare variants (defined here as MAF≤1%) which are not well-tagged by the single-nucleotide polymorphisms (SNPs) on genome-wide genotyping arrays are probably responsible for much of the heritability that remains unexplained^2^. Additionally, because purifying selection acts to remove deleterious alleles from the population, most variants that exert a sizable effect on complex traits, and that likely offer the best prospect for revealing biological mechanisms, should be particularly rare.

Rare variants are unevenly distributed between populations and difficult to represent effectively on commercial genotyping arrays, as evidenced by relatively sparse association findings even from large array-based studies of coding variants^3–6^. Discovering rare variant associations will therefore almost certainly require exome or genome sequencing of very large numbers of individuals. However, the sample size required to reliably identify rare-variant associations remains uncertain; most sequencing studies to date have identified few novel associations, and theoretical analyses confirm that they have been underpowered to do so^7^. These analyses also suggest that power to detect rare variant associations varies enormously between populations that have expanded in isolation from recent bottlenecks compared to those that have not.

In isolated populations that expand rapidly following a bottleneck, alleles that pass through the bottleneck often rise to a much higher frequency than in other populations^8–10^. If the bottleneck was recent, even deleterious alleles under negative selection may remain relatively frequent in these populations, resulting in increased power to detect association with disease-related traits. The Finnish population exemplifies this type of history. It grew from bottlenecks occurring 2,000-4,000 years ago in the founding of the early-settlement regions of southern and western Finland; internal migration in the 15^th^ and 16^th^ centuries to the late-settlement regions of northern and eastern Finland created additional bottlenecks^11^. The subsequent rapid growth of the Finnish population (to ~5.5 million, larger than any other human isolate) generated sizable geographic sub-isolates in late-settlement regions.

Geneticists have long noted that the bottlenecks that were so prominent in Finland’s recent history caused 36 Mendelian disorders to be much more common in Finland than in other European countries, while several other disorders are much less common, a phenomenon termed “the Finnish Disease Heritage”^12^. The identification of mutations for 35 of these disorders has confirmed that they mostly concentrate in late settlement regions^12^. Additional studies demonstrated, in these regions, an overall enrichment of deleterious variants more extreme compared to other isolates or to Finland generally^13–15^
. We reasoned that this enrichment would enable exome sequencing studies of late-settlement Finland to be better powered than studies in other populations to systematically investigate the impact of low-frequency variants on disease-related quantitative traits. Based on this expectation, we formed such a sample (“FinMetSeq”) from two Finnish population-based cohort studies: FINRISK and METSIM (see Methods).

Using >1.4 M variants identified and genotyped by successful exome sequencing of 19,292 FinMetSeq participants, we conducted single-variant association analysis with 64 clinically relevant quantitative traits^16,17^. We identified 43 novel associations with deleterious variants in 25 traits: 19 associations (11 traits) in FinMetSeq and 24 associations (20 traits) in a combined analysis of FinMetSeq with an additional 24,776 Finns from three cohorts for which imputed array-based genome-wide genotype data were available. Nineteen of the 26 variants underlying these 43 novel associations were unique to Finland or enriched >20-fold in FinMetSeq compared to non-Finnish Europeans (NFE).

We demonstrate that (1) a well-powered exome sequencing study can identify numerous rare alleles, each of which has a substantial effect on one or more traits in the individuals who carry them, and (2) exome sequencing in a population that has expanded after recent population bottlenecks is an extraordinarily efficient strategy to discover such effects. As most of the novel putatively deleterious trait-associated variants that we identified are unique to or highly enriched in Finland, similarly powered studies of these variants in non-Finnish populations might require hundreds of thousands or even millions of participants. Additionally, the geographical clustering of these enriched alleles, like the Finnish Disease Heritage mutations, demonstrates that the distribution of trait-associated rare alleles may vary significantly between locales within a country.

## RESULTS

### Genetic variation

We attempted to sequence the protein-coding regions of 23,585 genes covering 39 MB of genomic sequence in 20,316 FinMetSeq participants. After extensive quality control, we included in downstream analysis 19,292 individuals sequenced to 47x mean depth (Methods, **Supplementary Table 1**). We identified 1,318,781 single nucleotide variants (SNVs) and 92,776 insertion/deletion (indel) variants, with a mean of 20,989 SNVs and 604 indel variants per individual. The majority (87.5%) of SNVs identified were rare (MAF<1%); 40.5% were singletons (**Table 1**). Each participant carried 15 singleton variants on average, 17 rare (MAF≤1%) protein truncating variants (PTVs; annotated as stop gain, essential splice site, start loss, or frameshift) alleles, and 171 common (MAF>1%) PTVs (**Supplementary Table 2**). Frameshift indels accounted for the largest proportion of PTVs (31% of rare, 42% of common), while stop gain variants were the most frequent type of protein truncating SNVs (29% of rare, 20% of common).

**Table 1.**
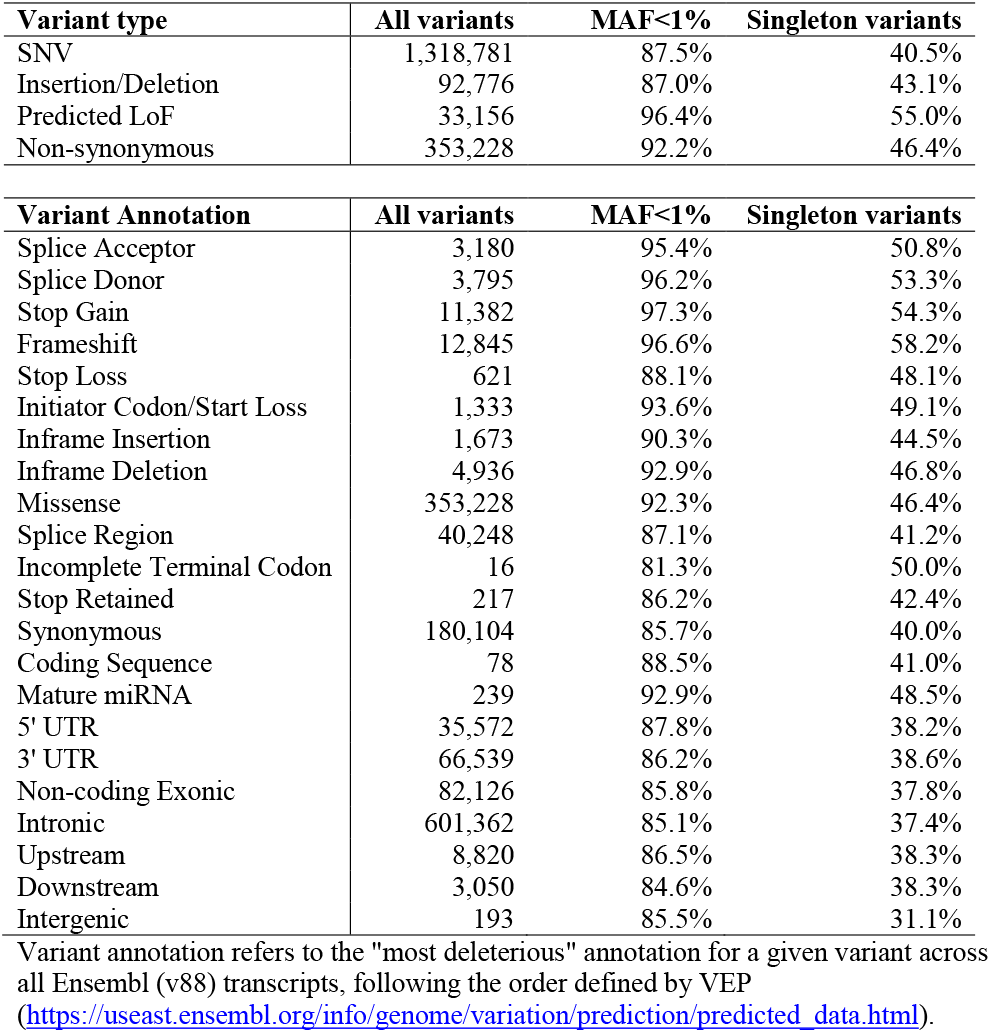
Sequence variants identified using whole-exome sequencing of 19,292 FinMetSeq participants. Percentages are the percent of all variants in the given category to either have MAF <1% or to be singleton variants.

| **Variant type** | **All variants** | **MAF<1%** | **Singleton variants** |
| --- | --- | --- | --- |
| SNV | 1,318,781 | 87.5% | 40.5% |
| Insertion/Deletion | 92,776 | 87.0% | 43.1% |
| Predicted LoF | 33,156 | 96.4% | 55.0% |
| Non-synonymous | 353,228 | 92.2% | 46.4% |

| **Variant Annotation** | **All variants** | **MAF<1%** | **Singleton variants** |
| --- | --- | --- | --- |
| Splice Acceptor | 3,180 | 95.4% | 50.8% |
| Splice Donor | 3,795 | 96.2% | 53.3% |
| Stop Gain | 11,382 | 97.3% | 54.3% |
| Frameshift | 12,845 | 96.6% | 58.2% |
| Stop Loss | 621 | 88.1% | 48.1% |
| Initiator Codon/Start Loss | 1,333 | 93.6% | 49.1% |
| Inframe Insertion | 1,673 | 90.3% | 44.5% |
| Inframe Deletion | 4,936 | 92.9% | 46.8% |
| Missense | 353,228 | 92.3% | 46.4% |
| Splice Region | 40,248 | 87.1% | 41.2% |
| Incomplete Terminal Codon | 16 | 81.3% | 50.0% |
| Stop Retained | 217 | 86.2% | 42.4% |
| Synonymous | 180,104 | 85.7% | 40.0% |
| Coding Sequence | 78 | 88.5% | 41.0% |
| Mature miRNA | 239 | 92.9% | 48.5% |
| 5' UTR | 35,572 | 87.8% | 38.2% |
| 3' UTR | 66,539 | 86.2% | 38.6% |
| Non-coding Exonic | 82,126 | 85.8% | 37.8% |
| Intronic | 601,362 | 85.1% | 37.4% |
| Upstream | 8,820 | 86.5% | 38.3% |
| Downstream | 3,050 | 84.6% | 38.3% |
| Intergenic | 193 | 85.5% | 31.1% |

Variant annotation refers to the "most deleterious" annotation for a given variant across all Ensembl (v88) transcripts, following the order defined by VEP (https://useast.ensembl.org/info/genome/variation/prediction/predicted_data.html).

We compared variant allele frequencies in FinMetSeq to those of NFE control exomes from the Genome Aggregation Database (gnomAD v2.1, **Extended Data Fig. 1**). As in previous smaller-scale comparisons of Finnish and NFE exomes, in FinMetSeq we observe a depletion of the rarest alleles (singletons and doubletons) and a relative excess of more common variants (minor allele count, MAC ≥5) compared to NFE for all classes of variants. This effect is particularly marked for alleles predicted to be deleterious (**Extended Data Fig. 2**).

### Single-variant association analyses

We tested for association between genetic variants in FinMetSeq and 64 clinically relevant quantitative traits measured in members of both FINRISK and METSIM (**Supplementary Table 3**). We adjusted lipid and blood pressure traits for lipid lowering and antihypertensive medication use, respectively, adjusted all traits for covariates using linear regression (**Supplementary Table 4**), and inverse normalized trait residuals to generate normally distributed traits for genetic association analysis that assumed an additive model (Methods). Based on common variants, 62 of 64 traits exhibited significant heritability (P<0.05; h^2^ range 5.0-52.5%; **Fig. 1A, Supplementary Table 5**), and there was substantial correlation between traits, phenotypically and genetically (**Fig. 1B**).

**Figure 1.**
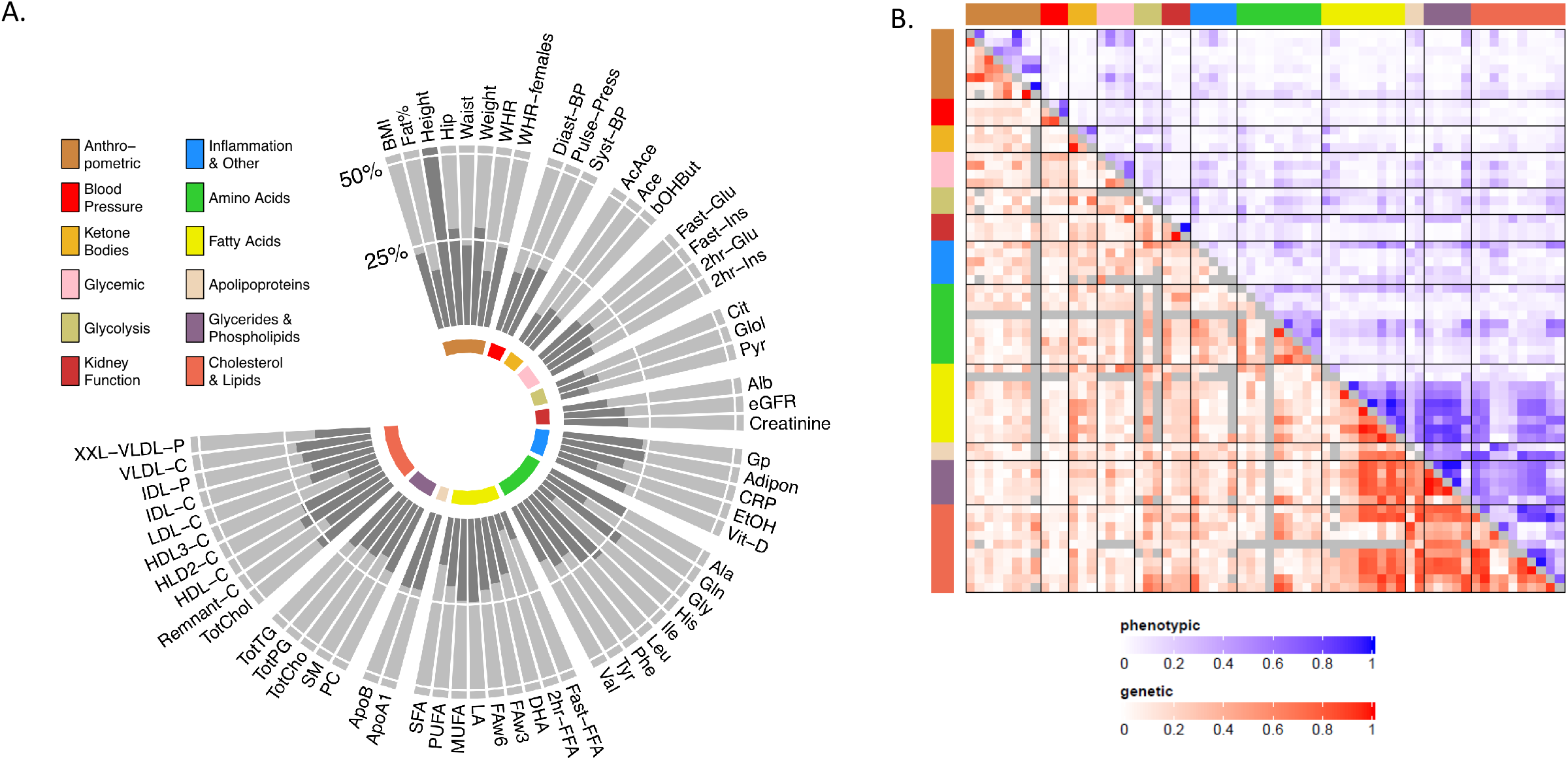
Characterization of traits by heritability, Pearson correlation, and genetic correlation. Traits in both figures are in the same order, clockwise in A, and left to right and top to bottom in B, and following the trait group color key. A) Estimated heritability (h_*x*_^2^) for each of the 64 traits included in association analysis. Heritability is based on ~205,000 common variants from GWAS arrays available in 13,342 unrelated individuals. Height has the highest heritability estimate at 52.5%. Estimates of trait heritability for metabolic measures are somewhat lower than previous reports (Kettunen, 2012) because estimates are from population-level data as opposed to twin studies and heritability was estimated from covariate adjusted and inverse normal transformed residuals, rather than raw trait values. Trait abbreviations are listed in Supplementary Table 3. All traits are significantly heritable except for 2hr-FFA (Fatty Acid) and His (Amino Acid), see Supplementary Table 5 for estimates, SEs, and P-values. B) Heatmap of: 1) absolute Pearson correlations of standardized trait values in upper triangle, and 2) absolute values of the genetic correlation, *ρ*_G_(x,y), in lower triangle, where *ρ*_G_(*x,y*) is the estimated genetic correlation of traits *x* and *y*. Values below the diagonal in gray had non-estimable genetic correlations.

We tested the 64 traits for single-variant associations with the 362,996 to 602,080 genetic variants with MAC ≥3 among the 3,558 to 19,291 individuals measured for each trait (**Supplementary Tables 3 & 4**). Association results are available for download and can be explored interactively with PheWeb (http://pheweb.sph.umich.edu/FinMetSeq/) and via the Type 2 Diabetes Knowledge Portal (www.type2diabetesgenetics.org). We identified 1,249 trait-variant associations (P<5×10^−7^) at 531 variants (**Supplementary Table 6**), with 53 of 64 traits associated with at least one variant (**Fig. 2A**). All 1,249 associations remained significant after multiple testing adjustment across the exome and across the 64 traits with a hierarchical procedure setting average FDR at 5% (Methods). Using the hierarchical FDR procedure, we detected an additional 287 trait-variant associations at these 531 variants (**Supplementary Table 7**). These additional associations reflect the high correlation between a subset of lipid traits, e.g. high-density lipoprotein cholesterol (HDL-C) and apolipoprotein A1 (ApoA1). Given the diversity of traits assessed in these cohorts, these associations may shed additional light on the biology of measures that have been less frequently assayed in large GWAS, such as intermediate density lipoproteins (IDL) and very-low-density lipoprotein (VLDL) particles. Of the 531 associated variants, 59 (11%) were rare (MAF≤1%); by annotation, 200 (38%) were coding, 108 (20%) missense, and 11 (2%) protein truncating. Furthermore, minor alleles at >10-fold increased frequency in FinMetSeq compared to NFE are substantially more likely to be associated with a trait compared to variants with similar or lower MAF in FinMetSeq compared to NFE (OR=4.92, P=2.6×10^−5^; **Extended Data Fig. 3**).

**Figure 2.**
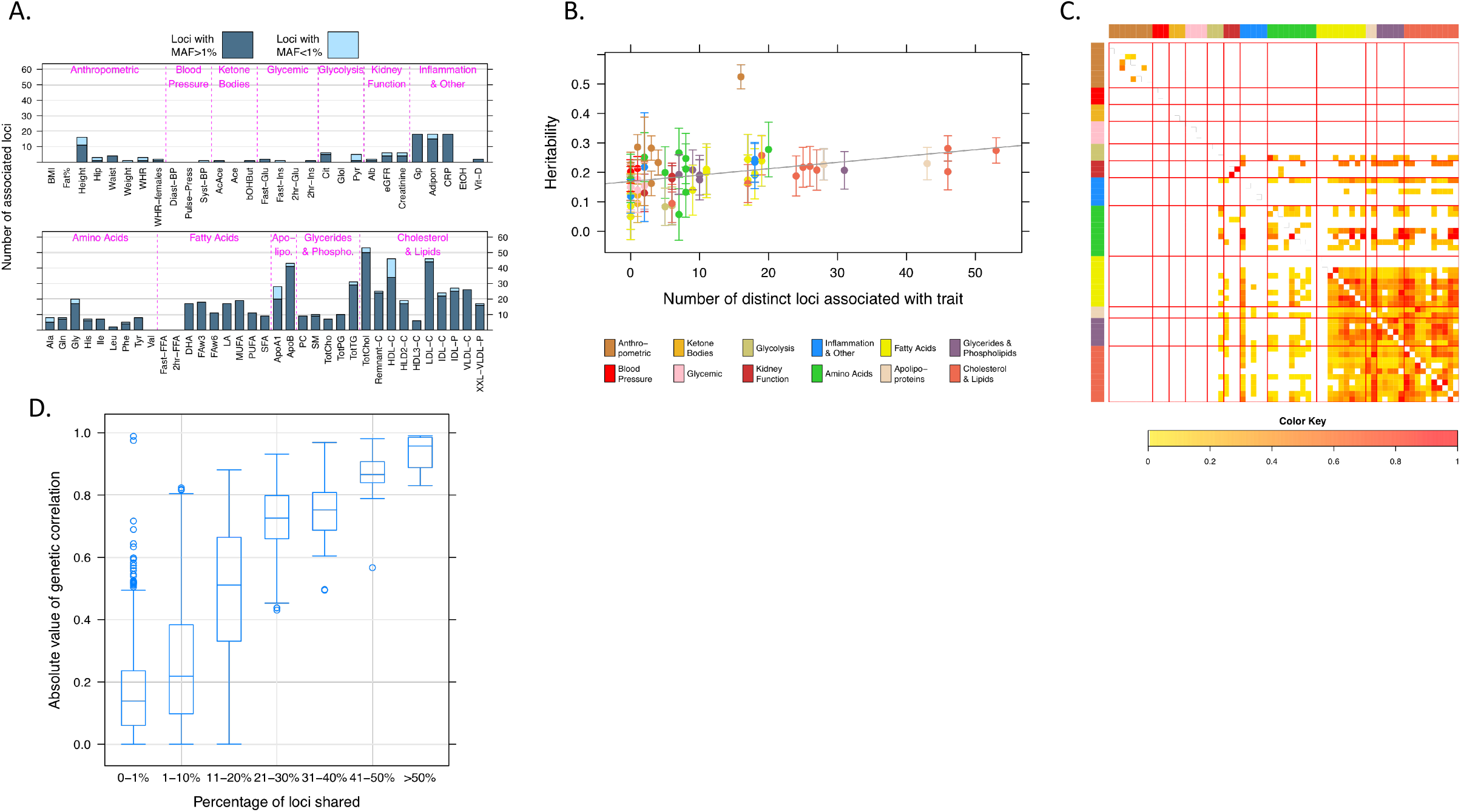
Characterization of discovered associations. A) Number of genomic loci associated with each trait. Each bar is subdivided into common (MAF>1%, dark blue) and rare (MAF<1%, light blue). Traits are sorted by group as in Figure 1. B) Relationship between estimated heritability and number of genomic loci detected for each trait. Each trait is colored by trait group following the trait group color key. Vertical bars indicated ±2 standard errors of the heritability estimate. The gray line shows the linear regression fit, shown to indicate the general trend. C) Heatmap of shared genomic associations by pairs of traits. For traits *x* and *y*, the color in row *x* and column *y* reflects the number of loci associated with both traits divided by the number of loci associated with trait *x*. Traits are presented in the same order as in 2A, and the side and top color bars reflect the trait groups. D) Relationship between estimated genetic correlation and extent of sharing of genetic associations. For each pair of traits, the extent of locus sharing is defined as the number of loci associated with both traits divided by the total number of loci associated with either trait. The bar within each box is the median, the box represents the inter-quartile range, whiskers extend up to 1.5x the interquartile range, and outliers are presented as individual points. Analysis using the absolute value of the Pearson correlation of the residual series results in a very similar pattern.

We clumped associated variants within 1 Mb and with r^2^>0.5 into a single locus, irrespective of the associated traits (Methods). After clumping, the 531 associated variants represented 262 distinct loci (597 trait-locus pairs, **Supplementary Table 6**); 158 of the 262 loci (60%) consisted of a single trait-associated variant. As expected, the number of associated loci per trait was positively correlated with trait heritability (r=0.38, P=8.8×10^−4^). Height was a noticeable outlier, with relatively few associations considering its high estimated heritability (**Fig. 2B**).

The majority of variants and loci (61%) were associated with a single trait; 4% were associated with ≥10 traits. Overlapping associations (**Fig. 2C**) strongly reflect the relationships exhibited by both trait and genetic correlations (**Fig. 1B**). For example, rs113298164, a missense variant in *LIPC* (p.Thr405Met), is associated with 11 traits, including cholesterols, fatty acids, apolipoproteins, and cholines. Similarly, the estimated genetic correlation of trait pairs is a strong predictor of the probability for a trait pair to share associated loci (**Fig. 2D**).

To determine which of the 1,249 single-variant associations were distinct from known GWAS associations for the same traits, we repeated association analysis for each trait conditional on published associated variants (P<10^−7^) for the corresponding trait in the EBI GWAS Catalog (December 2016 release). Of the 1,249 trait-variant associations, 478 (at 213 of 531 variants) remained significant (P<5×10^−7^) after conditional analysis, representing 126 of the original 262 loci, including at least one conditionally significant locus for each of 48 traits (**Supplementary Table 8**). The conditionally-associated variants were more often rare (24% vs. 11%), more likely to alter or truncate the resulting protein (31% vs. 22%), and more frequently >10x enriched in FinMetSeq relative to NFE (19% vs. 10%) compared to the full set of associated variants.

### Gene-based association analyses

To identify genes associated with the 64 traits, we performed aggregate tests of protein coding variants, grouping variants using three different masks. Mask 1 comprised PTVs of any frequency; Masks 2 and 3 also included missense variants with MAF<0.1% or 0.5% predicted to be deleterious by five algorithms (Methods). We identified 54 gene-based associations with P<3.88×10^−6^ (adjusting for testing a maximum of 12,890 genes containing at least two qualifying variants) and with multi-trait FDR<0.05, analogous to the threshold used for single-variant association testing (Methods). Fifteen of these associations required ≥2 variants to achieve significance (i.e. the association was not driven by a single strongly associated variant; **Supplementary Table 9**). Extremely rare (MAC≤3) PTVs drove the association of eight traits with *APOB* (**Extended Data Fig. 4**). We found a novel association between two very rare stop gain variants in *SECTM1* and HDL2 cholesterol (P=7.2×10^−7^, **Extended Data Fig. 5**). *SECTM1* encodes an interferon-induced transmembrane protein that is negatively regulated by bacterial lipopolysaccharide (LPS)^18^. The association could reflect the role of HDL particles in binding and neutralizing LPS in infections and sepsis^19^.

### Replication and follow-up of single-variant associations in three additional Finnish cohorts: Identification of novel coding, deleterious variant associations

We attempted to replicate the 478 single-variant associations from FinMetSeq (unconditional and conditional P≤5×10^−7^) and to follow-up the 2,120 suggestive but sub-threshold associations from FinMetSeq (unconditional 5×10^−7^<P≤5×10^−5^, conditional P≤5×10^−5^) in 24,776 participants from three Finnish cohort studies for which varying subsets of the 64 FinMetSeq traits were available: FINRISK^20,21^ participants not sequenced in FinMetSeq (n=18,215), the Northern Finland Birth Cohort 1966^22^ (n=5,139), and the Helsinki Birth Cohort^23^ (n=1,412). For each of the three cohorts, we carried out genotype imputation using the Finnish-specific SISu v2 reference panel (http://www.sisuproject.fi), which is comprised of 5,380 haplotypes from whole-genome based sequencing and 10,184 haplotypes from whole-exome based sequencing in coding regions, and then used the same single-variant association analysis strategy employed in FinMetSeq. We then carried out meta-analysis of the three imputation-based studies to test for replication of associated FinMetSeq variants (“replication analysis”) and four-study meta-analysis with FinMetSeq to follow-up suggestive associations (“combined analysis”; Methods).

We obtained data for 448 of the 478 significant variant-trait associations (191 of the 213 requested variants). Of the 448 associations for which we had replication data, 439 (98.0%) had the same direction of effect in replication analysis as in FinMetSeq; 392 of the 448 replicated at P<0.05 (87.5%; **Supplementary Table 10**). We also obtained data to follow up 1,417 of the 2,120 sub-threshold associations (1,014 of the 1,554 requested variants); >60% of the variants that we could not follow up were very rare in FinMetSeq and were not present in the SISu reference panel. Of the 1,417 sub-threshold trait-variant associations, 431 reached P<5×10^−7^ in the combined analysis (**Supplementary Table 11**).

Among the significant results from FinMetSeq or combined analysis, 43 associations were with 26 predicted deleterious variants that conditional analysis and literature review suggest are novel (**Table 2**). Nineteen such associations, at 15 deleterious coding variants, were significant in FinMetSeq (**Table 2; Supplementary Table 10**). Twelve of these associations replicated (P<0.05) in the replication analysis and remained significant in the combined analysis; for the other seven associations we either did not have replication data (six associations) or did not replicate but had very low power (<5%) in the replication analysis (one association). Four of the 15 variants were PTVs; 11 were missense variants predicted to be deleterious by at least one of five prediction algorithms. Another 24 associations, with 16 variants (two PTVs and 14 missense variants predicted to be deleterious), only reached significance in the combined analysis (**Table 2; Supplementary Table 11**). Five variants with significant associations in FinMetSeq alone were associated with additional traits in combined analysis (**Table 2**).

**Table 2.**
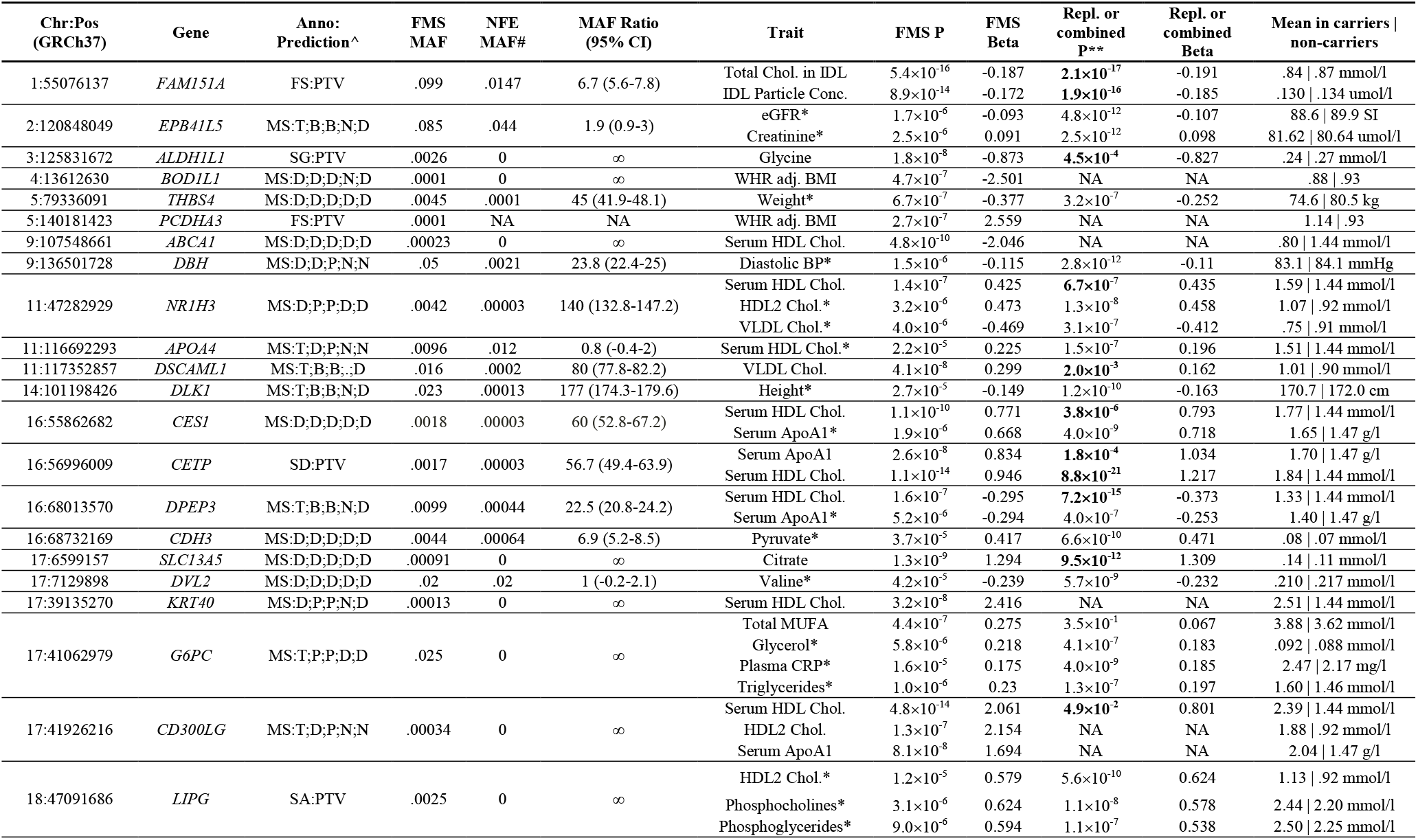
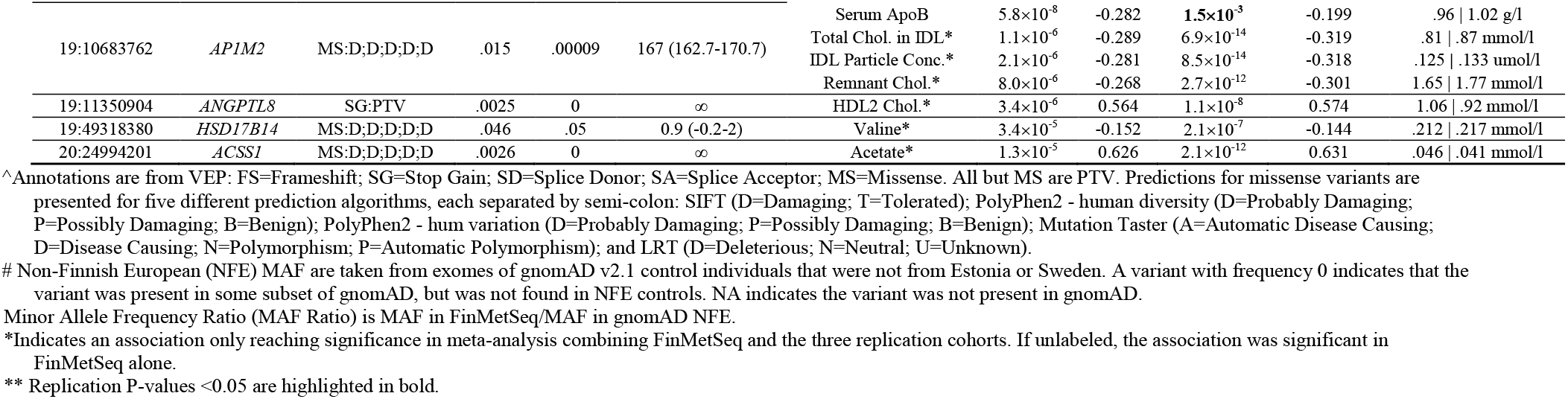
Associations with predicted deleterious variants that conditional analysis and literature review suggest are novel. These associations reach exome-wide significance in FinMetSeq alone or in a combined analysis of FinMetSeq with three replication cohorts.

Of the 43 associations shown in **Table 2**, 34 were with 19 variants either seen only in Finland or enriched by >20-fold in FinMetSeq compared to NFE (13 of 15 variants in FinMetSeq and 11 of 16 variants in combined analysis with five variants overlapping). Identifying associations for these 19 variants would have required much larger samples in NFE populations than in FinMetSeq (**Fig. 3A & B**). We provide brief summaries relating each of these highly enriched associations to known biology and prior genetic evidence relating to the respective genes in **Supplementary Information**. We highlight a few of the most striking findings, below.

**Figure 3.**
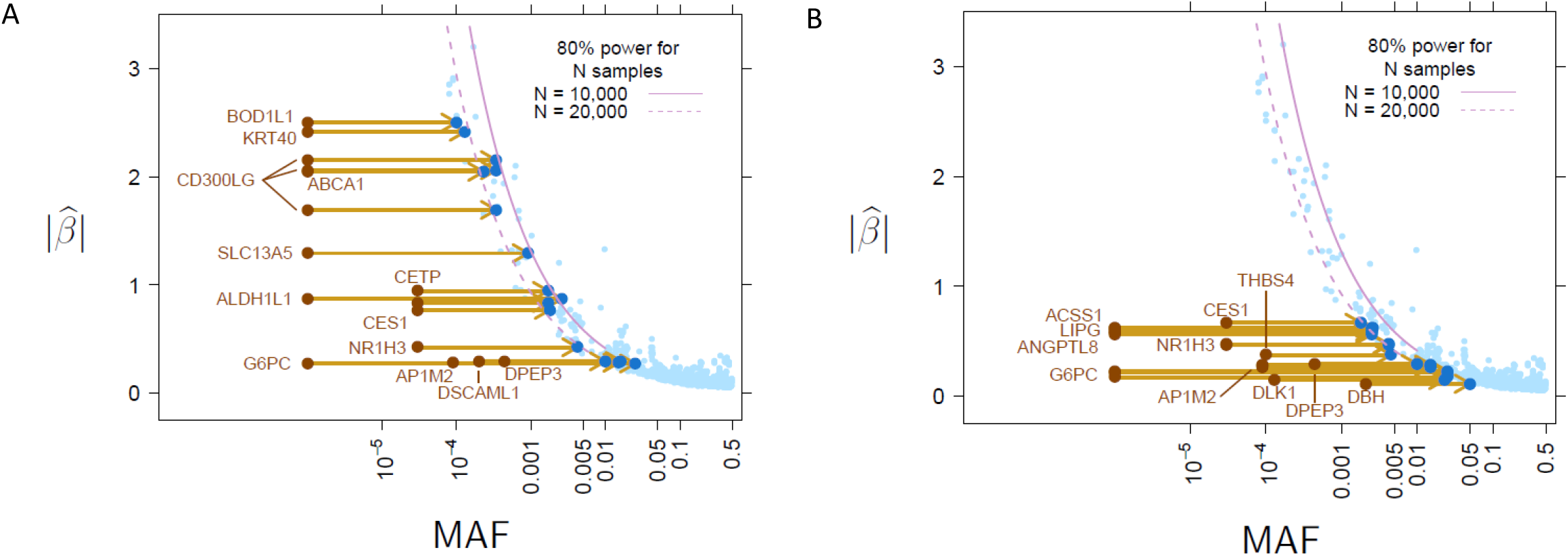
Allelic enrichment in the Finnish population and its effect on genetic discovery. A) Relationship between MAF and estimated effect size for associations discovered in FinMetSeq exomes alone. Each variant reaching significance in FinMetSeq is plotted. Those associations highlighted in Table 2 are represented with a dark blue point (FinMetSeq MAF) and a corresponding brown point reflecting the NFE MAF (gnomAD). The purple lines indicate the 80% power curves for significance at 5×10^−7^ for sample sizes of 10,000 and 20,000. The right end of the power curve for N=20,000 terminates at MAF = 0.007. Plots show the dramatic increase in power due to higher relative frequency in Finland. B) Relationship between MAF and estimated effect size for associations discovered in the combined analysis. Same plot as in A, highlighting the variants in Table 2 only reaching significance in the combined analysis.

#### Anthropometric traits

As a group these are among the most extensively investigated quantitative traits, with thousands of common variant associations reported, most of very small effect^24–28^. We identified several rare, large effect variants for these traits, including a predicted damaging missense variant (rs200373343, p.Arg94Cys) in *THBS4* 45X more frequent in FinMetSeq than in NFE and associated in the combined analysis with a mean decrease in body weight of 5.9 kg (**Table 2**). *THBS4* encodes thrombospondin 4, a matricellular protein found in blood vessel walls and highly expressed in heart and adipose tissue^29^. *THBS4* is involved in local signaling in the developing and adult nervous system, and may function in regulating vascular inflammation^30^. Coding variants in *THBS4* and other thrombospondin genes have been implicated in increased risk for heart disease^31–33^.

We identified a predicted damaging missense variant (rs2273607, p.Val104Met) in *DLK1* that is 177X more frequent in FinMetSeq than in NFE and is associated in the combined analysis with a mean decrease in height of 1.3 cm (**Table 2**). *DLK1* encodes Delta-Like Notch Ligand 1, an epidermal growth factor that interacts with fibronectin and inhibits adipocyte differentiation. Uniparental disomy of *DLK1* causes Temple Syndrome and Kagami-Ogata Syndrome, characterized by pre- and postnatal growth restriction, hypotonia, joint laxity, motor delay, and early onset of puberty^34–36^. Paternally-inherited common variants near *DLK1* have been associated with child and adolescent obesity, type 1 diabetes, age at menarche, and central precocious puberty in girls^37–39^. Homozygous null mutations in the mouse ortholog Dlk-1 lead to embryos with reduced size, skeletal length, and lean mass^40^, while in Darwin’s finches, SNVs at this locus have a strong effect on beak size^41^.

#### HDL-C

Two novel variants with large effects on HDL-C in FinMetSeq are absent in NFE. The predicted deleterious missense variant rs750623950 (p.Arg112Trp) in *CD300LG* is associated in FinMetSeq with a mean increase in HDL-C of 0.95 mmol/l, and also associated with HDL2-C and ApoA1 (**Table 2**). *CD300LG* encodes a type I cell surface glycoprotein. A missense variant in *ABCA1* (rs765246726, p.Cys2107Arg) is associated in FinMetSeq with a mean reduction in HDL-C of 0.64 mmol/l (**Table 2**). Fifteen more variants (including ten which are absent in NFE) contributed to a strong *ABCA1* gene-based association signal (P=2.2×10^−13^; **Supplementary Table 9, Extended Data Fig. 6**). *ABCA1* encodes the cholesterol efflux regulatory protein, which regulates cholesterol and phospholipid metabolism. Individuals who are homozygotes or compound heterozygotes for any of several *ABCA1* mutations produce very little HDL-C and experience the manifestations of severe hypercholesterolemia.

#### Amino Acids

A stop gain variant (rs780671030, p.Arg722X) in *ALDH1L1* is associated in FinMetSeq with a mean reduction in serum glycine levels of 0.03 mmol/l but is not observed in NFE (**Table 2**); this effect may increase risk for several cardiometabolic disorders^42,43^. *ALDH1L1* encodes 10-formyltetrahydrofolate dehydrogenase, which competes with the enzyme serine hydroxymethyltransferase to alter the ratio of serine to glycine in the cytosol. Although rs780671030 was the strongest associated variant, gene-based association tests suggest that additional PTVs and missense variants in *ALDH1L1* also alter glycine levels (P=1.4×10^−20^, **Extended Data Fig. 7, Supplementary Table 9**).

#### Ketone bodies

A predicted damaging missense variant (rs201013770, p.Phe517Ser) in *ACSS1* is associated in the combined analysis with mean increased serum acetate level of 0.005 mmol/l but is not observed in NFE (**Table 2**). *ACSS1* encodes an acyl-coenzyme A synthetase and plays a role in the conversion of acetate to acetyl-CoA. In rodents, increased acetate levels lead to obesity, insulin resistance, and metabolic syndrome, mediated by activation of the parasympathetic nervous system^44^.

### Associated variants and disease endpoints

Newly available GWAS data from the FinnGen project^45^ enabled us to test the hypothesis that deleterious variants responsible for our novel quantitative trait associations (**Table 2**) could also contribute to disease endpoints related to these traits. FinnGen has particularly rich data on such endpoints as the samples are largely drawn from Finnish hospital biobanks. In total, we examined 22 disease endpoint phenotypes for all 25 available variants in **Table 2**. Three variants were associated with disease endpoints in FinnGen at a Bonferroni-corrected threshold of P<0.05/(22×25)=9.0×10^−5^ (**Supplementary Table 12**).

A predicted damaging missense variant (17:39135270:A/G; p.Ser32Pro) in *KRT40* which is not observed in NFE and associated in FinMetSeq with a mean elevation in HDL-C of 1.07 mmol/l (**Table 2**), is associated in FinnGen with increased risk for pancreatitis. While this is the first disease association reported for this gene, the type I keratin family, of which *KRT40* is a member, is believed to play an important role in regulating exocrine pancreas homoeostasis^46^. A 29 bp deletion on chromosome 1 causes a frameshift in *FAM151A* which is 6.7X more frequent in FinMetSeq than NFE and associated in FinMetSeq with both decreased total cholesterol in IDL and decreased IDL particle concentration (**Table 2**), is associated in FinnGen with decreased risk of myocardial infarction. The interpretation of this association is complicated by the fact that the variant is also present in an overlapping transcript (*ACOT11*), a gene that plays a role in fatty acid metabolism and lies <1 MB from a well-known cardioprotective variant in *PCSK9*. Finally, a predicted damaging missense variant (rs77273740; p.Arg65Trp) in *DBH* that is 23.8X more frequent in FinMetSeq than in NFE and is associated with a mean decrease of 1 mmHg in diastolic blood pressure in our combined analysis (**Table 2**), is associated in FinnGen with decreased risk for hypertension. Distinct loci in this gene have previously been shown with mean arterial pressure and this variant was included in a gene-based association with mean arterial pressure^5,6^.

### Replication outside of Finland: UK Biobank

To begin to assess the generalizability outside of Finland of the novel associations that we detected, we attempted to replicate associations from our combined Finnish analyses in the UK Biobank (UKBB), a European sample that is approximately ten-fold larger. Across eight anthropometric and blood pressure traits for which UKBB data are publicly available, our Finnish combined analysis had identified 31 trait-variant associations reaching P<5×10^−7^. More than a quarter of these variants (8 of 31) were not present in the UKBB database. Of the remaining 23 associations, 20 were to variants that were common in FinMetSeq (MAF> 1%) and had a comparable frequency in UKBB; 15 (75%) of these variants showed association in UKBB at P<0.05/23=2.2×10^−3^ (Bonferroni correction for 23 tests). Of the three rare variants in this analysis, all of which were enriched at >10x frequency in FinMetSeq compared to UKBB, none showed association in UKBB (**Supplementary Table 13**). Even after adjusting for winner’s curse^47^ and with a sample size of 340,000-360,000, we had <50% power to detect all three of these associations in UKBB (**Supplementary Table 13**). This comparison supports the argument that extremely large samples will be needed in most other populations to achieve the power for rare variant association studies that we have observed in Finland.

### Geographical clustering of associated variants

Given the concentration within sub-regions of northern and eastern Finland of most Finnish Disease Heritage mutations^48^, we hypothesized that the distribution of rare trait-associated variants discovered through FinMetSeq might also display geographical clustering. In support of this hypothesis, principal component analysis revealed broad-scale population structure within the late-settlement region among 14,874 unrelated FinMetSeq participants whose parental birthplaces are known (**Extended Data Fig. 8**). Consistent with our hypothesis, parental birthplaces were significantly more geographically clustered for carriers of PTVs and missense alleles than for carriers of synonymous alleles, even after adjusting for MAC (**Supplementary Tables 14A, 14B**).

To enable finer scale analysis of the distribution of variants within late-settlement Finland, we delineated geographically distinct population clusters using patterns of haplotype sharing among 2,644 unrelated individuals with both parents known to be born in the same municipality (Methods, **Extended Data Fig. 9**). Taking the cluster that is most genetically similar to early-settlement Finland as a reference, we compared variant counts for different functional classes and frequencies between this reference cluster and each of the other 12 clusters that contained ≥100 individuals (**Fig. 4, Supplementary Tables 15, 16**). In the two clusters that represent the most heavily bottlenecked late-settlement regions (Lapland and Northern Ostrobothnia), we observed a marked deficit of singletons and significant enrichment of variants at intermediate frequency compared to other clusters. This pattern is most significant for missense variants, which are present in the exome in large numbers; PTVs show consistently greater enrichment, but with less statistical significance likely due to very small counts (**Fig. 4**). Two clusters in which we observed marked enrichment of singletons, Surrendered Karelia and South Ostrobothnia, showed the highest levels of genetic diversity across the frequency spectrum, likely reflecting relatively recent gene flow into these regions from neighboring countries (Russia and Sweden, respectively, **Fig. 4**).

**Figure 4.**
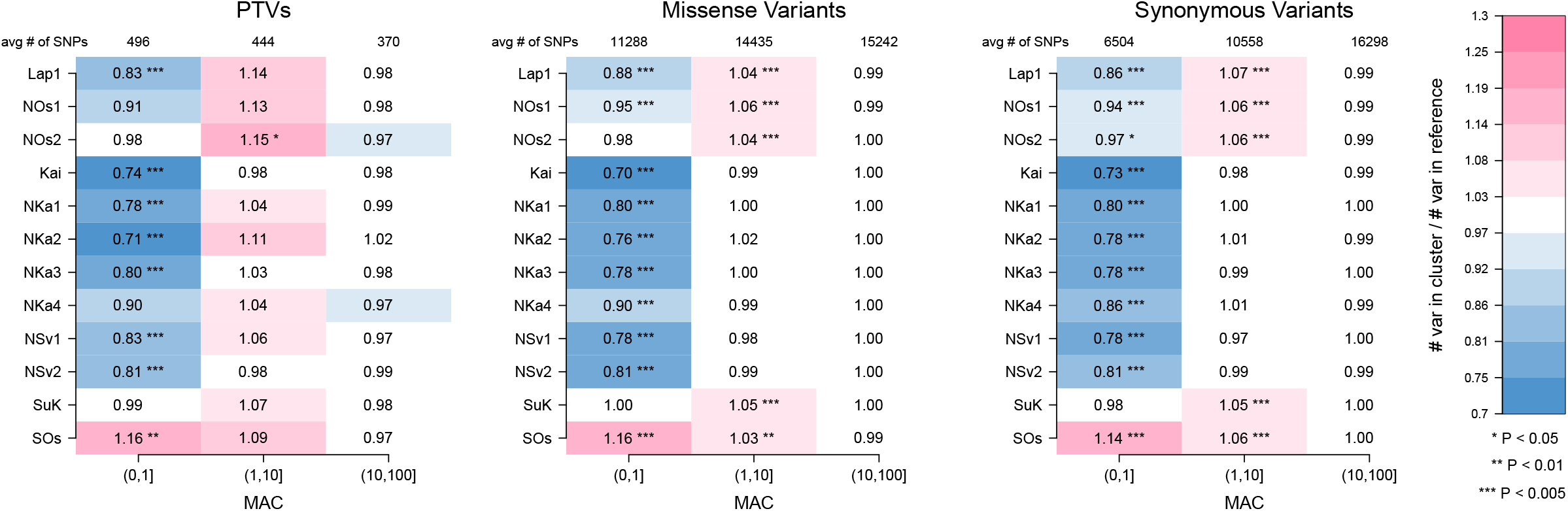
Regional variation in allele frequencies by functional annotation. Enrichment of functional allelic class in sub-populations (regions) of Northern and Eastern Finland. For each minor allele count bin, we computed the ratio of number of variants found in each subpopulation to an internal reference subpopulation (NSv3), after down-sampling the frequency spectra of all populations to 200 chromosomes. Pink cells represent an enrichment (ratio >1), blue cells represent a depletion (ratio <1). The 12 sub-populations with sample size >100 are shown. The results are consistent with multiple independent bottlenecks followed by subsequent drift in Northern and Eastern Finland, particularly for populations in Lapland and Northern Ostrobothnia. Abbreviations for regions: Kainuu (Kai), Lapland (Lap1, Lap2), Northern Karelia (NKa1, NKa2, NKa3, NKa4), Northern Ostrobothnia (NOs1, NOs2, NOs3, NOs4), Northern Savonia (NSv1, NSv2, NSv3), Southern Ostrobothnia (SOs), and Surrendered Karelia (SuK). For more detailed information on region definitions see Supplementary Table 15. Confidence intervals on the enrichment ratios, and their P-values, are presented in Supplementary Table 16.

We observed particularly strong geographical clustering among variants >10X enriched in FinMetSeq compared to NFE (**Fig. 5A, Extended Data Fig. 10, Supplementary Table 17**). We further characterized geographical clustering for FinMetSeq-enriched trait-associated variants, by comparing the mean distances between birthplaces for parents of minor allele carriers to those of non-carriers (**Supplementary Table 18**). Most such variants were highly localized. For example, for variant rs780671030 in *ALDH1L1*, which may be unique to Finns, the mean distance between parental birthplaces is 135 km for carriers and 250 km for non-carriers (P<1.0×10^−7^, **Fig. 5B**). In contrast, few of the variants that displayed sub-threshold association in FinMetSeq but that showed significant associations in the combined analysis were significantly geographically clustered within Finland (**Supplementary Table 18**).

**Figure 5.**
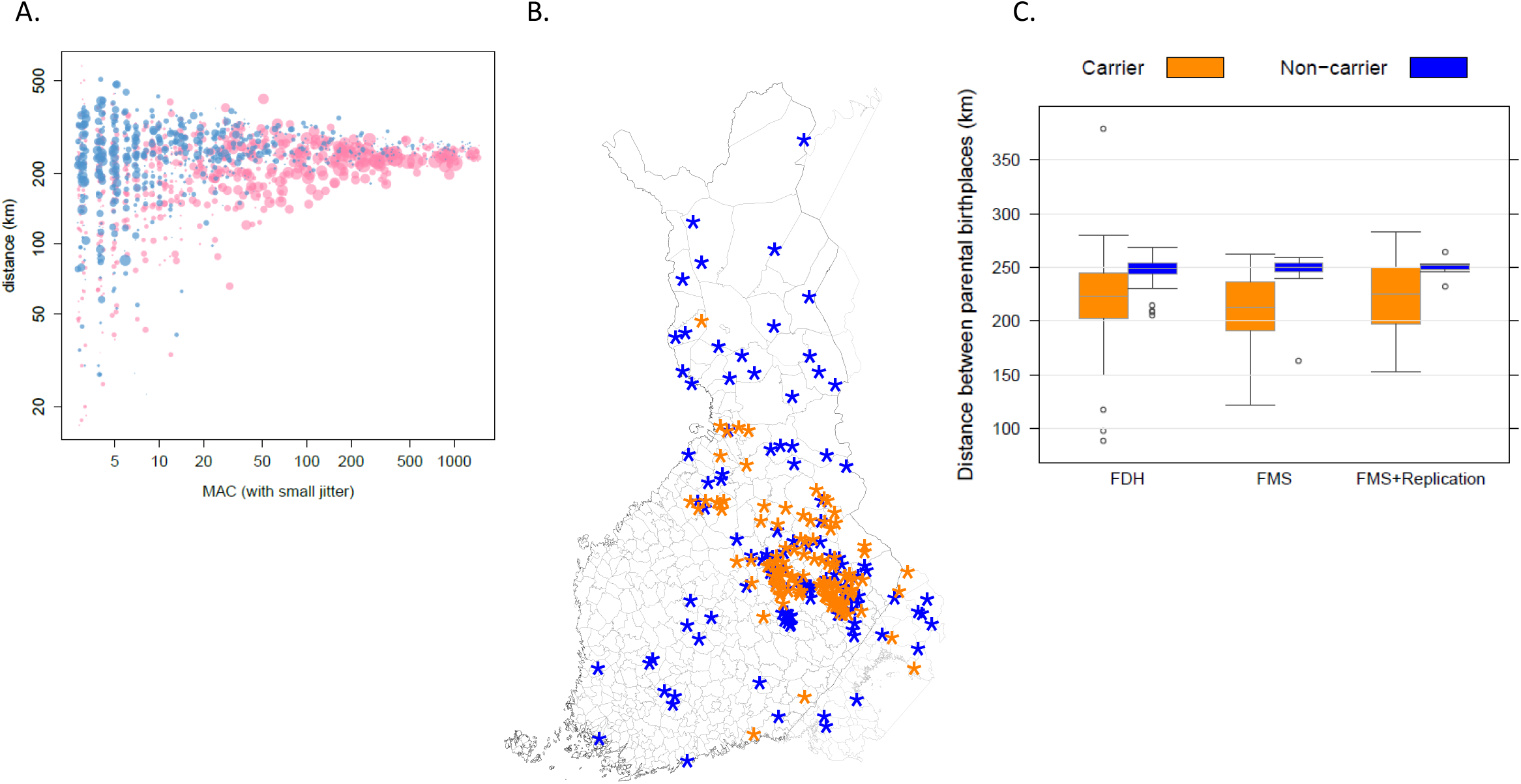
Geographical clustering of associated variants. A) Geographical clustering of PTVs as a function of MAC and frequency enrichment over NFE from gnomAD. For each PTV (r^2^≤0.02, MAC≥3, MAF≤0.05) we computed the mean distance between birth places of available parents of all carriers of the minor allele. We compared the frequency of the minor allele in FinMetSeq to gnomAD NFE. Blue and pink colors denote the frequency is lower or higher in FinMetSeq than in gnomAD NFE, respectively. The size of the point is proportional to the logarithm of the frequency ratio difference. In general, we observe that variants enriched in FinMetSeq are more geographically clustered. B) Example of geographical clustering for a trait associated variant. The birth locations of all parents of carriers (orange) and a matching number of parents of non-carriers (blue) of the minor allele for variant chr3:125831672 (rs780671030, p.Arg722X) in *ALDH1L1* are displayed on a map of Finland. This variant is associated with serum glycine levels in FinMetSeq and has a frequency of 0 in NFE samples from gnomAD. The parents of carriers are born on average 135 km apart, the parents of non-carriers on average 250 km apart (P<10^−7^ by permutation). C) Comparison of geographical clustering between Finnish Disease Heritage (FDH) mutations and trait-associated variants that are >10x more frequent in FinMetSeq than in NFE. The degree of geographical clustering (based on parental birthplace) is comparable between carriers of those variants that showed significant associations in FinMetSeq alone (FMS) and carriers of FDH mutations, and greater than that seen in carriers of variants that showed significant association only in the combined analysis (FMS+Replication). For all variants, carriers of the minor allele displayed greater clustering than non-carriers. The bar within each box is the median, the box represents the inter-quartile range, whiskers extend up to 1.5× the interquartile range, and outliers are presented as individual points.

Finally, we compared the geographic clustering of FinMetSeq-enriched trait-associated variants to that of 35 Finnish Disease Heritage mutations carried by ≥3 FinMetSeq individuals with known parental birthplaces. FinMetSeq carriers of monogenic Finnish Disease Heritage mutations and FinMetSeq carriers of trait-associated variants identified in FinMetSeq displayed a comparable degree of geographic clustering. This clustering was dramatically greater than that observed for the non-carriers of both sets of variants (**Fig. 5C**), suggesting that rare variants associated with complex traits may be much more unevenly distributed geographically than has been appreciated to date.

## DISCUSSION

We have demonstrated that a well-powered exome sequencing study of deeply phenotyped individuals can identify numerous rare variants associated with medically relevant quantitative traits. The variants that we identified may provide a useful starting point for studies aimed at uncovering biological mechanisms or fostering clinical translation. For example, further investigation of the p.Arg722X variant in *ALDH1L1* associated with reduced serum glycine could help elucidate the role of this gene in astrocyte function, a topic of growing interest in neurobiology. Glycine is a key inhibitory neurotransmitter localized to astrocytes^49^, while the mouse ortholog, *Aldh1l1*, is the primary marker for astrocytes in experimental research, since it is strongly expressed in astrocytes, but not in neurons^50^.

The substantial power of this study for discovering rare variant associations derives from the occurrence, in the recently expanded and heavily bottlenecked populations of northern and eastern Finland, of a large pool of deleterious variants that appear unique to Finland or at frequencies orders of magnitude greater than in NFE. This observation motivates a strategy for scaling up the discovery of rare variant associations by prioritizing the sequencing of populations beyond Finland that have expanded in isolation from recent bottlenecks. Examples of other such populations include those of Ashkenazim^51^, Iceland^52^, Quebec^53^, highland regions of Latin America^54^, and geographically isolated regions of larger European countries such as Sardinia^55^ and Crete^10^. In each of these populations, genetic drift has almost certainly caused a different set of alleles to pass through the corresponding population-specific bottlenecks, enriching some variants while depleting others. The numerous rare-variant associations that could be identified by sequencing available, phenotyped samples across multiple population isolates could rapidly increase our understanding of the genetic architecture of complex traits. One caveat is that the extended LD blocks that are typical in such populations may make it difficult to identify the causative variant from among multiple deleterious variants within an association region^56^.

Recent studies have suggested a continuity between the genetic architectures of complex traits and disorders classically considered monogenic^57,58^. Our results offer strong support for this continuity, not only in identifying numerous deleterious variants with large effects on quantitative traits, but in demonstrating that such variants show geographical clustering comparable to that of the mutations responsible for the Finnish Disease Heritage.

The use of a Finland-specific genotype reference panel^59^ to impute FinMetSeq variants into array-genotyped samples from three other Finnish cohorts enabled us to identify many additional novel associations. This result suggests that using sequence data from a subset of individuals in each population to impute variants in array-genotyped samples from the same population is a cost-effective strategy for detecting rare-variant associations. However, the clustering in FinMetSeq of deleterious trait-associated variants within limited geographical regions and our inability to follow-up >700 sub-threshold associations from FinMetSeq for which the associated variants were not present in the Finnish imputation reference panel, emphasize the importance of extensively representing regional subpopulations when designing such reference panels, to account for fine-scale population structure.

To fully realize the value of large-scale sequencing studies in population isolates, it will be necessary to increase the richness of phenotypes available in sequenced cohorts from these populations. For example, we associated <100 of the >24,000 deleterious, highly enriched variants identified in FinMetSeq with one of the 64 cardiometabolic-related quantitative traits studied here. In Finland, the national health care system and the population’s willingness to participate in biomedical research mean that extensive medical records and population registries are available for mining additional phenotype data, and create an opportunity for callback by genotype for further phenotyping and collection of biological samples^60^. Notably, the associations we identified to disease endpoints in FinnGen give a hint of the discoveries that will be possible when that database reaches its full size of 500,000 participants. The insights gained from such efforts will accelerate the implementation of precision health, informing projects in larger, more heterogeneous populations which are still at an early stage^61^.

## Acknowledgements

Thanks to Terri Teshiba for coordinating ethical permissions and samples. Thanks to Sini Kerminen, Daniel Lawson, and George Busby for discussions and providing scripts to run fineSTRUCTURE. SR was supported by the Academy of Finland Center of Excellence in Complex Disease Genetics (Grant No 312062), Academy of Finland (No. 285380), the Finnish Foundation for Cardiovascular Research, the Sigrid Juselius Foundation, Biocentrum Helsinki and University of Helsinki HiLIFE Fellow grant. VR acknowledges support by RFBR, research project No. 18-04-00789 A. VS was supported by the Finnish Foundation for Cardiovascular Research. CS and LS received funding from HG006695, HL113315, and MH105578. MAK is supported by a Senior Research Fellowship from the National Health and Medical Research Council (NHMRC) of Australia (APP1158958). He also works in a unit that is supported by the University of Bristol and UK Medical Research Council (MC_UU_12013/1). The Baker Institute is supported in part by the Victorian Government’s Operational Infrastructure Support Program. AUJ, DR, LJS, HMS, RW, PY, XY, and MB received funding from DK062370. SKS, CWKC, and NBF received funding from HL113315 and NS062691. The METSIM study was supported by grants from Academy of Finland (No. 321428), the Sigrid Juselius Foundation, the Finnish Foundation for Cardiovascular Research, Kuopio University Hospital, and Centre of Excellence of Cardiovascular and Metabolic Diseases supported by the Academy of Finland (ML). Sequencing was funded by 5U54HG003079, and AEL, KMS, HJB, CCC, CJK, KLK, DCK, DEL, JN, TJN, SKD, NOS, IMH, and RKW were funded by 5U54HG003079 and 5UM1HG008853-03.

## Author Contributions

AEL, LJS, RKW, AaP, VS, ML, SR, MB, and NBF designed the study. AEL, KMS, HJA, RSF, DCK, DEL, JN, TJN, and JV produced and quality-controlled the sequence data. AEL, AUJ, ArP, HMS, MAK, VS, and ML produced and quality-controlled the clinical data. AEL, KMS, CWKC, SKS, ASH, LS, MP, CCC, AUJ, CJK, KK, VR, DR, JV, RW, PY, and XY analyzed data. JGE, MAK, MRJ, and MM provided replication data. HL, SKD, NOS, IMH, CS, SR, MB, and NBF supervised experiments and analyses. AEL, KMS, CWKC, SKS, CS, MB and NBF wrote the paper. AEL, KMS, CWKC, and SKS contributed equally to this work. NBF and MB jointly supervised this work.

## Competing interests statements

VS has participated in a conference trip sponsored by Novo Nordisk and received a honorarium from the same source for participating in an advisory board meeting. He also has ongoing research collaboration with Bayer Ltd.

HL is a member of the Nordic Expert group unconditionally supported by Gedeon Richter Nordics and has received an honorarium from Orion.

Correspondence and requests for materials should be addressed to.

## Data Availability

The sequence data can be accessed through dbGaP using the following study numbers: FINRISK: phs000756, METSIM: phs000752. Association results can be accessed at http://pheweb.sph.umich.edu/FinMetSeq/.

## METHODS

### METSIM and FINRISK studies: designs, phenotypes, and sequenced participants

**METSIM** is a single-site study comprised of 10,197 men randomly selected from the population register of Kuopio, Eastern Finland, aged 45 to 73 years at initial examination from 2005 to 2010^17,62^. The goal of METSIM is to investigate genetic and non-genetic factors associated with Type 2 Diabetes (T2D), cardiovascular disease (CVD), insulin resistance, and related traits. The METSIM study protocol includes collection of data on CVD history and risk factors, measurements of height, weight, waist, hip, blood pressure, and collection of a blood sample for laboratory measurements and DNA extraction. Diagnoses of myocardial infarction^63^, stroke^64^, and peripheral vascular disease were verified based on medical records at baseline. We attempted exome sequencing of all METSIM study participants.

**FINRISK** is a series of health examination surveys carried out by the National Institute for Health and Welfare (formerly National Public Health Institute) of Finland every five years beginning in 1972^65^. The surveys are based on random population samples from five (six in 2002) geographical regions of Finland. Participants were selected by 10-year age group, sex, and study area. Survey sample sizes have varied from 7,000 to 13,000 individuals and participation rates from 60% to 90%. The age-range was 25 to 64 years until 1992 and 25 to 74 years since 1997. The survey includes a self-administered questionnaire, a standardized clinical examination carried out by specifically trained study nurses, and collection of a blood sample for laboratory measurements and DNA extraction^66^. For exome sequencing, we chose 10,192 participants from the 1992, 1997, 2002, and 2007 FINRISK surveys from northeastern Finland (former provinces of North Karelia, Oulu, and Lapland). This selection was based on the hypothesis that the rapid growth in isolation of the populations of this region from severe bottlenecks in the 16^th^-17^th^ centuries might have resulted in deleterious variants rising to a much higher frequency than in other populations.

METSIM participants fasted for more than 10 hours prior to blood draw. FINRISK participants were instructed to fast for four hours before the scheduled examination and to avoid heavy meals earlier in the day; duration of fasting was recorded. Laboratory measurements in METSIM included an oral glucose tolerance test with samples at 0, 30, and 120 minutes (only fasting measurements in known diabetics) for glucose, insulin, proinsulin, and free fatty acids, as well as fasting laboratory measurements including lipids, lipoproteins, apolipoproteins, adiponectin, and hs-CRP. Most of these measurements were carried out in FINRISK non-fasting samples; two-hour oral glucose tolerance tests with glucose and insulin measured at 0 and 120 minutes were carried out in subsets of FINRISK 1992, 2002 and 2007 participants. Both studies include proton NMR metabolomics measurements of lipoprotein subclasses, their lipid concentrations and composition, apolipoprotein A-I and B, multiple cholesterol and triglyceride measures, albumin, various fatty acids, and numerous low-molecular-weight metabolites, including amino acids, glycolysis related measures and ketone bodies^67,68^.

METSIM and FINRISK participants are followed up regularly via record linkage using the Finnish health registries, allowing for near complete follow-up of multiple disease diagnoses; participants may also be called back in person for additional studies. Participants in both studies provided informed consent, and all study protocols were approved by the Ethics Committees at the participating institutions (FINRISK cohorts 1992 & 1997: National Public Health Institute of Finland; FINRISK 2002, 2007, & 2012: Ethical Review Board of the Hospital District of Helsinki and Uusimaa; METSIM: Research Ethics Committee, Hospital District of Northern Savo IRB #1).

### Selection of traits, harmonization, exclusions, covariate adjustment, and transformation

Of the 257 quantitative metabolic and cardiovascular traits measured in both METSIM and FINRISK, we selected 64 traits for association analysis in the current study based on clinical relevance for cardiovascular and metabolic health (**Supplementary Tables 3, 4**).

#### Exclusions

We excluded 126 individuals, 92 with type 1 diabetes and 34 women who were pregnant at the time of phenotyping, from all analyses, and 3,088 individuals with T2D from analyses of glycemic traits. For traits influenced by food consumption (amino acids, fatty acids, LDL cholesterol, total triglycerides, and glycemic traits), we excluded individuals not fasting for at least 8 hours after their last meal. A complete list of exclusions can be found in **Supplementary Table 4**.

#### Trait adjustments

For individuals on antihypertensive medication at the time of testing, we added 15 and 10 mmHg to the measured values of systolic and diastolic blood pressures, respectively^69,70^. For individuals on lipid regulating medications, we multiplied the measured total cholesterol by 1.25^71^. For FINRISK participants, we calculated LDL cholesterol (LDL-C) levels using the Friedewald equation (LDL-C = adjusted total cholesterol – HDL-C – (triglycerides/2.2)) and excluded individuals with elevated triglycerides (>2.5mmol/l); LDL-C was measured directly in METSIM participants and for participants on lipid regulating medication, values were divided by 0.7^72^. All trait adjustments are listed in **Supplementary Table 4**.

#### Trait transformations and adjustment for covariates

We prepared quantitative traits for association analysis separately for METSIM and FINRISK participants by linear regression on trait-specific covariates; skewed variables were log transformed prior to linear regression analysis. Both before and after log transformation, we examined all variables for normality and for outliers. Log transformation was adequate in all cases to approximate normality for initial covariate adjustment. Outliers, of which there were no more than 5 individuals with values >5SD for any trait prior to adjustment and inverse normalization, were not removed. Covariates for these regression analyses always included covariates age and age^2^ for METSIM and sex, age, age^2^, and cohort year for FINRISK. Trait transformations and trait-specific covariates are listed in **Supplementary Table 4**. Several traits were adjusted for sex hormone treatment, which included women on contraceptives or hormone replacement therapy. We transformed residuals from these initial regression analyses to normality using inverse normal scores.

### Exome sequencing

We carried out exome sequencing in two phases.

<u>Phase 1</u> We quantified the 10,379 Phase 1 DNA samples with the Quant-iT PicoGreen dsDNA reagent on a Varioskan Microplate Reader (ThermoFisher Scientific) and randomly parsed samples with adequate DNA (>250ng) into cohort specific files. We then re-arrayed samples using the BioMicroLab XL20 (USA Scientific) to ensure equal numbers of METSIM and FINRISK samples on each 96-well plate, alternating samples between studies in consecutive positions within and across plates, to reduce opportunities for between-study batch effects.

We constructed dual indexed libraries using 100-250ng of genomic DNA and the KAPA HTP Library Kit (KAPA Biosystems) on the SciClone NGS instrument (Perkin Elmer). The target insert size was 250 bp. We then pooled twelve libraries prior to hybridization with the SeqCap EZ HGSC VCRome (Roche) reagent that targets 45.1 Mb (23,585 genes and 189,028 non-overlapping exons) of the human genome. Each library pool contained samples from both cohorts and was comprised of 300-400 ng of each individual library for a total library input of 3.6-4.8 μg into the initial hybridization. We estimated the concentration of each captured library pool by qPCR (Kapa Biosystems) to produce appropriate cluster counts for the HiSeq2000 platform (Illumina). We then generated 2×100bp paired-end sequence data yielding ~6 Gb per sample to achieve a coverage depth of ≥20x for ≥70% of targeted bases for every sample.

<u>Phase 2</u> We quantified, prepared, pooled, and captured the 9,937 Phase 2 samples just as in Phase 1. Here we generated 2×125 bp paired-end sequencing reads on the HiSeq2500 1T to again achieve a coverage depth of ≥20x for ≥70% of targeted bases for every sample.

#### Contamination detection, sequence alignment, sample QC, and variant calling

We aligned sequence reads to human genome reference build 37 using bwa-mem (v0.7.7), marked duplicates with picard MarkDuplicates (v1.113; http://broadinstitute.github.io/picard), and realigned indels with the GATK^73^ IndelRealigner (v2.4). We used BamUtil clipOverlap (v1.0.11; http://genome.sph.umich.edu/wiki/BamUtil:_clipOverlap) to mark overlapping bases from paired reads resulting from short insert fragments.

For each sample, we required ≥70% of target bases sequenced at ≥20x depth, and SNV genotype array concordance >90% if SNV array data were available. We used verifyBamID^74^ (v1.1.1) to detect and exclude samples with estimated contamination >3%. Where available, we also used existing genotype data with verifyBamID to detect and exclude sample swaps. Of 20,316 samples attempted, 13 failed sequencing, 35 were sample swaps, 760 either had low genotype concordance, sex mismatch, or estimated contamination >3%, and four had discrepancies between reported and genotype-estimated relationships (**Supplementary Table 1**).

We called SNVs and short indels using the recommended best practices for cohort-level calling with GATK^73^ (v3.3). For each individual, we called bases and identified variant sites for all targeted exome bases and 500 bp of sequence up and downstream of each target region using HaplotypeCaller, resulting in calling substantial numbers of non-exonic variants. We merged these calls in batches of 200 individuals using CombineGVCFs and recalled genotypes for all individuals at all variable sites with GenotypeGVCFs.

After merging genotypes for the 19,378 samples that passed preliminary QC checks, we filtered SNVs and indels separately using the recommended best practices for Variant Quality Score Recalibration (VQSR). We used the set of true positive variants provided in the GATK resource bundle (v2.5; build37) for training the VQSR model after restricting to sites in targeted exome regions. After assessment with VQSR, we retained variants for which we identified ≥99% of true positive sites used in the training model (i.e. truth sensitivity) for both SNVs and indels.

Following initial variant filtering, we decomposed multi-allelic variants into bi-allelic variants, left-aligned indels, and dropped redundant variants using vt^75^ (version 0.5). We filtered variants with >2% missing calls and/or Hardy-Weinberg p-value<10^−6^. We applied an additional filter removing variants with an overall allele balance (AB; alternate AC/sum of total AC) <30% in genotyped samples. We then excluded 86 individuals with >2% missing variant calls yielding a final analysis set of 19,292 individuals.

### Array genotypes, genotype imputation, and integrated exome+imputation panel

METSIM participants were previously genotyped with the Illumina OmniExpress array; genotyping and quality control are described elsewhere^76^. FINRISK participants were previously genotyped in several batches on different arrays^21^. We lacked genotype array data for 1,488 participants (57 METSIM, 1,431 FINRISK). From the available genotype array data, we generated three datasets: 1) a merged array-based genotype call set of all variants present in ≥90% of array-genotyped individuals across both cohorts; 2) a merged Haplotype Reference Consortium (HRC) v1.1 imputed data set of the array-based genotypes; 3) an integrated data set containing genotyped and imputed array-based variants and exome sequence variants (HRC+exome). Where there was overlap between the sequence and imputed genotypes, we kept the sequence-based genotypes. We excluded the 1,488 individuals with no array data from the HRC+exome panel.

We prepared array genotypes for imputation using the Imputation Preparation and Checking tool (http://www.well.ox.ac.uk/~wrayner/tools/HRC-1000G-check-bim.v4.2.5.zip) and used the Michigan Imputation Server^77^ (www.imputationserver.sph.umich.edu) to impute genotypes using the HRC (v1.1) reference panel^78^. METSIM samples were imputed in a single batch. FINRISK samples were imputed in batches based on the genotyping array and/or center where genotypes were generated.

### Annotation

We annotated the final set of variants passing QC using Ensembl’s variant effect predictor (VEP v76)^79^ and Combined Annotation-Dependent Depletion^80^ (CADD v1.2). We employed five *in silico* algorithms implemented in dbNSFP v2.4 (https://sites.google.com/site/jpopgen/dbNSFP) to predict the functional impact of missense variants: PolyPhen2 HumDiv and HumVar^81^, LRT^82^, MutationTaster^83^, and SIFT^84^.

### Association testing

#### Single variants

We carried out single-variant association tests for transformed trait residuals with genotype dosages for variants with MAC≥3 assuming an additive genetic model. We used the EMMAX^85^ linear mixed model approach, as implemented in EPACTS (v3.3.0; http://genome.sph.umich.edu/wiki/EPACTS), to account for relatedness between individuals. We used genotypes for sequenced variants with MAF≥1% to construct the genetic relationship matrix (GRM).

#### Conditioning on associated variants from prior GWAS

To differentiate association signals identified in this study from known association signals, for each trait we performed exome-wide association analysis conditioning on variants previously associated (P<10^−7^) with that trait. We compiled a list of known variants for each trait from the EBI GWAS catalog (https://www.ebi.ac.uk/gwas/downloads; December 4, 2016 version), from recent papers, and from manuscripts in preparation of which we were aware^76,86–88^. The keywords from the GWAS catalog we used to assign known variants to each of our traits are listed in **Supplementary Table 19**. We also manually curated the associations from Inouye et al.^89^ and Kettunen et al.^86^ to attribute “blood metabolite” associations to the specific associated metabolites.

Using the combined HRC+exome panel (see above), we pruned each trait-specific list of associated variants (“GWAS variants”) based on linkage disequilibrium (LD) (r^2^>0.95). Twenty-three GWAS variants were not present in the HRC+exome panel. For 17 of these 23 variants, we identified a proxy (r^2^>0.80) variant instead; we excluded the remaining six variants from conditional analysis. The list of variants included in conditional analysis for each trait is included in **Supplementary Table 20**. We extracted genotypes for variants used in conditional analysis from the integrated HRC+exome panel and converted dosages to alternate allele counts by rounding to the nearest integer (0, 1, or 2). We imputed missing genotypes for the 1,488 individuals without array data using the mean genotype dosage for purposes of conditional analysis.

For conditional analysis for each trait, we ran association analysis using the same linear mixed model approach as in unconditional analysis but including the complete set of pruned GWAS variants as covariates in the association test. Following conditional association, we further determined novelty based on absence of the variant from OMIM descriptions, ClinVar, and extensive literature searches.

#### Defining loci

The set of >1.4M variants tested for association includes variants in LD. To identify the number of distinct associations for each trait, we performed LD clumping using Swiss (https://github.com/welchr/swiss). We selected the subset of variants with (1) unconditional P<5×10^−7^ or (2) both unconditional and conditional P<5×10^−5^ for at least one trait. For each variant in this subset, we provided Swiss with the minimum unconditional p-value across all traits. The clumping procedure starts with the variant with the smallest p-value (index variant), and merges into one locus all variants within ±1 Mb that have r^2^>0.5 with the index variant. The procedure is repeated iteratively until no variants remain. Trait associations with variants in the same locus are considered to represent the same signal and trait associations with variants in different loci to be distinct signals.

#### Calculating effects and variance explained of individual variants

For novel variants highlighted in **Table 2** we evaluated the effect of each variant on the trait values. We did this by calculating the mean trait value in carriers and non-carriers, assuming no homozygous carriers. Differences noted are the difference in the two means.

Given that the effect estimates from our association tests are standardized, we calculated variance explained for a given variant with the equation 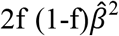, where f is the minor allele frequency and 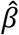 is the estimated effect size. The variance explained is included in **Supplementary Table 8**.

#### Gene-based testing

We carried out gene-based association tests using the mixed model implementation of SKAT-O^90^, which tests for the optimal mixture of burden and dispersion-style multi-marker tests while adjusting for relatedness between individuals using the same GRM calculated for the single-variant tests. EMMAX and the mixed model version of SKAT-O (mmskat) are implemented in EPACTS.

We implemented gene-based tests using three different, but nested, sets of variants (variant “masks”):

1. PTVs at any allele frequency with VEP annotations: frameshift_variant, initiator_codon_variant, splice_acceptor_variant, splice_donor_variant, stop_lost, stop_gained;
2. PTVs included in (1) plus missense variants with MAF<0.1% scored as “damaging” or “deleterious” by all five functional prediction algorithms;
3. PTVs included in (1) plus missense variants with MAF<0.5% scored as “damaging” or “deleterious” by all five functional prediction algorithms.

For each trait and mask, we only tested genes with at least two qualifying variants. Each mask contained a different number of genes with at least two qualifying variants: up to 7,996, 12,795, and 12,890 for the three masks, respectively. The exact number of genes tested varied by trait due to sample size. We first used a Bonferroni-corrected exome-wide threshold for 12,890 genes, which corresponds to a threshold of P<3.88×10^−6^. Analogous to single-variant association, we passed genes meeting this association threshold forward for additional consideration with hierarchical FDR correction as described below.

### Hierarchical FDR correction for testing multiple traits and variants

In controlling for multiple testing our goal was to make sure that, by looking across 64 traits, we did not increase the proportion of falsely discovered variants. To accomplish this, we adopted a FDR controlling procedure described in Peterson et al.^91^, which uses a hierarchical strategy to increase power while controlling type I error (**Supplementary Methods**). This procedure has two stages. Stage 1 identifies the set of R variants that are associated with at least one trait, controlling genome-wide FDR across all variants at 0.05. Stage 2 identifies all the traits that are associated with the discovered variants in a manner that guarantees an average FDR<0.05.

In Stage 1 we restricted ourselves to the R=531 variants that have an unconditional association P<5×10^−7^ with at least one trait. For these, we calculated a p-value for the hypothesis of no association between the variant and any of the 64 traits using Simes’ rule^92^, a combination rule that is robust to dependence between phenotypes. To account for the fact that we did an initial selection of these R variants from the total number of variants tested (T), we passed the Simes p-values to a Benjamini-Hochberg (BH) procedure that controls FDR at target level 0.05×R/T, a modification^93^ which guarantees that the FDR in the set of S variants discovered to be associated with at least one trait is less than 0.05.

In stage 2, to determine which traits are associated with the set of the S selected variants we apply the Benjamini and Bogomolov^94^ procedure. This procedure applies a multiplicity correction variant by variant, passing the 64 trait association p-values from each of the S selected variants and all 64 traits to a BH procedure that controls FDR at target level 0.05×S/T.

We applied this hierarchical correction to two different sets of results: the set of single-variant unconditional results and the set of gene-based test results. The gene-based tests used a threshold of P<3.88×10^−6^ to identify the R nominally significant genes in the first stage of the hierarchical procedure.

### Genotype validation

We validated exome sequence-based genotype calls using Sanger sequencing for METSIM carriers of 13 trait-associated very rare variants with MAF<0.1% in seven genes. All but one of 108 (99.1%) non-reference genotypes validated were concordant.

### Association replication in additional Finnish cohorts

We performed replication analysis of significant single-variant associations (P<5×10^−7^) and follow-up analysis of suggestive single-variant associations (P<5×10^−5^) in up to 24,776 individuals from three GWAS cohort studies: Northern Finland Birth Cohort 1966 (NFBC1966), the Helsinki Birth Cohort Study (HBCS), and FINRISK study participants not included in the exome sequencing portion of FinMetSeq.

A detailed description of the NFBC1966 study has been published previously and additional information is available at: http://www.oulu.fi/nfbc/node/18091^22^. The data used here, including clinical measurements and blood samples for genetic and NMR metabolite analyses, were collected at the 31-year follow-up in 1997. NFBC1966 samples (n=5,139) were genotyped on the Illumina 370k array.

The HBCS includes participants born in Helsinki from 1934-1944 and has been described elsewhere^23^; a basic description is available at https://thl.fi/fi/web/thlfi-en/research-and-expertwork/projects-and-programmes/helsinki-birth-cohort-study-hbcs-idefix. HBCS samples (n=1,412) were genotyped on the Illumina 610k array.

The FINRISK cohort was described in detail above, and participants (replication n=18,125) were genotyped in several batches on the Illumina 610k, CoreExome, or OmniExpress arrays^20,21^.

For each replication cohort, prior to phasing we performed genotype quality control batch-wise using standard quality thresholds for both sample-wise and variant-wise filtering: call rate>95%, HWE>10^−6^, MAF>5%. We pre-phased array genotypes with Eagle^95^ (v2.3) and imputed genotypes genome-wide with IMPUTE^96^ (v2.3.1) using the SISu v2 reference panel consisting of 2,690 sequenced Finnish genomes and 5,092 sequenced Finnish exomes. Following imputation, we assessed imputation quality by confirming sex, comparing sample allele frequencies with reference population estimates, and examining imputation quality (INFO score) distributions. We excluded any variant with INFO<0.7 within a given batch from all replication/follow-up analyses.

For each of the three cohorts, we matched, harmonized, covariate adjusted, and transformed available phenotypes as described above for FinMetSeq. We used the same covariates as for FINRISK. For each cohort, we ran single-variant association using the EMMAX linear mixed model implemented in EPACTS after generating kinship matrices from LD-pruned (command: plink --indep-pairwise 50 5 0.2) directly genotyped variants with MAF>5%.

### Association to disease endpoints in FinnGen

From a list of >1,100 disease endpoints available for analysis in the FinnGen project, we selected 22 we considered most likely to be related to the quantitative traits analyzed in FinMetSeq. As described in detail in Tabassum et al.^45^, variant associations with disease endpoints in FinnGen biobank participants were tested using SPAtest R package and adjusting for age, sex, and first 10 PCs in up to ~97,000 individuals.

### Association replication in UK Biobank

For the eight traits analyzed in FinMetSeq that were also available in the current UKBB release (height, weight, BMI, hip circumference, waist circumference, fat percentage, systolic blood pressure, and diastolic blood pressure), we extracted trait-variant association statistics for variants reaching P<5×10^−7^ in the FinMetSeq combined analysis from the analysis of unrelated white British individuals generated by the Neale lab (http://www.nealelab.is/uk-biobank). Seven of the eight traits had at least one associated variant and 23 of the total of 31 variants were available in UKBB. A comparison of association results is in **Supplementary Table 13**.

### Population genetic analyses

#### Identifying unrelated individuals

To identify a set of nearly independent common autosomal SNVs, we removed SNVs with MAF<5% and pruned the remaining SNVs in windows of 50 SNVs, in steps of 5 SNVs, such that no pair of SNVs had r^2^>0.2. We used the resulting 26,036 SNVs to estimate pairwise relationships among the 19,292 exome-sequenced individuals using KING^97^. We then removed one individual from each of the 4,418 pairs inferred by KING to have a relationship of 3rd degree or closer, resulting in a set of 14,874 (nearly) unrelated individuals for population genetic analyses.

#### Identifying sub-population clusters in FinMetSeq

We first combined exome sequence variants and a genome-wide set of 220,798 SNVs from GWAS arrays to provide a genome-wide backbone to aid in phasing and computing haplotype sharing. After removing variants with MAC<3, variants in known regions of long range LD^98^ and variants with HWE<10^−4^, we phased the remaining 764,696 variants using SHAPEIT^99^ (version 2, r837). To assess the substructure in our dataset while minimizing the effect of mixing due to recent population mobility, we focused on the 2,644 unrelated individuals born by 1955 whose parents were both born in the same municipality (irrespective of the birth location of the individual).

We identified sub-populations of the 2,644 individuals using ChromoPainter (version 2) and fineSTRUCTURE^100^ (version 2.0.8). We first used ChromoPainter to generate a pairwise co-ancestry matrix, which represents each individual’s DNA as a count of haplotype blocks copied from every other individual in the dataset. Following previous practices^101^, for computational efficiency, we estimated and fixed the switch and global emission rates as input for ChromoPainter on a subset of the data; cluster inference is known to be robust to up to 10-fold deviations of the estimated switch and emission rates^102^. For further computational speedup, we generated an initial clustering by applying a normal mixture model clustering^103^ (mclust package in R, version 5.1) to the top ten principal components of the coancestry matrix and used this initial cluster solution as seed to the fineSTRUCTURE analysis. We conducted 1 million Markov chain Monte Carlo (MCMC) iterations retaining one sample for every 1,000 iterations after discarding 3 million iterations as burn-in. After MCMC, we used fineSTRUCTURE to perform *post-hoc* refinement of cluster membership; we started with the MCMC sample with the highest posterior probability and reassigned membership, taking into account the cluster membership at each of the recorded MCMC samples^102^.

In total, we ran five MCMC chains using fineSTRUCTURE, retaining the configuration with highest posterior probability for further analysis. We confirmed convergence of the fineSTRUCTURE MCMC runs by calculating Geweke’s convergence diagnostic using the coda package (version 0.18) in R to compare the number of inferred clusters in the first 10% and last 50% of the MCMC chain, and visual inspections of the general consistency of cluster memberships between independent MCMC chains. In total, we inferred 245 sub-population clusters among the 2,644 individuals.

We inspected the initial clustering solution from fineSTRUCTURE by examining for each individual the estimated proportion of their haplotype length derived from each of the inferred clusters using non-negative least squares^102,104^. This approach showed many individuals derived a substantial proportion of their haplotype length not from the cluster initially assigned by fineSTRUCTURE, but instead from a different but related sub-cluster on the fineSTRUCTURE hierarchical clustering tree, suggesting redundancy in fineSTRUCTURE-inferred clusters. We therefore combined related clusters by successively merging pairs of clusters that resulted in the smallest decrease in the posterior probability of the fineSTRUCTURE hierarchical clustering tree. At each merge, we reorganized individuals into merged cluster memberships and re-estimated the haplotype-sharing profile for each individual. We iteratively merged the hierarchical tree until ≥90% of individuals were assigned to the cluster where they also derive the highest proportion of haplotype sharing, resulting in 16 clusters for the 2,644 reference individuals, each named based on the most common parental birthplaces of its members (**Supplementary Table 15**).

#### Enrichment of predicted functionally deleterious alleles in Finland

We assessed enrichment of predicted functionally deleterious alleles in Finland by comparing the 14,874 nearly unrelated (pairwise kinship coefficient <0.0448) FinMetSeq individuals to the 14,944 NFE control exomes in gnomAD, excluding from the NFE individuals from the neighboring countries of Estonia and Sweden in which substantial numbers of Finns reside. We analyzed sites with base quality score >10, mapping quality score >20, and coverage equal to or greater than that found in ≥80% of variable sites (17.73X in FinMetSeq, 32.27X in gnomAD), resulting in ~38.6 Mbp for comparisons. We considered only the two most common alleles at each site. We contrasted the proportional site frequency spectra for FinMetSeq and NFE for five functional variant categories (PTVs, missense, synonymous, UTR, and intronic variants) after accounting for sample size differences between datasets by down-sampling both datasets to 18,000 chromosomes.

We also assessed the enrichment of functional alleles within subpopulations of the FinMetSeq dataset. Of the 16 sub-population clusters identified by fineSTRUCTURE, we used as the reference population a cluster for which the highest proportion of the parents of its members were from the southwestern, “early-settlement” part of Finland (NSv3, **Supplementary Table 15**). Twelve of the remaining 15 clusters also have >100 members and were used in subsequent analyses (**Supplementary Table 15**). We then compared the ratio of the site frequency spectra to the reference for PTVs, missense, and synonymous variants, again down-sampling both datasets to 200 haploid chromosomes to account for sample size differences. For a given comparison, we computed statistical evidence for enrichment or depletion at a given allele count bin by exact binomial test against a null of equal number of variants found in both the test and reference cluster.

#### Geographical clustering of predicted functionally deleterious alleles

We first generated a distance matrix tabulating the pairwise geographical distance in kilometers between the birthplaces of all available parents of unrelated sequenced individuals. For each variant of interest, we computed for the minor allele carriers in FinMetSeq the mean distance among all parent pairs. For example, for a variant with three carriers with information for five (of the possible six) parents, we computed the mean for all (5-choose-2 = 10) distances. We evaluated statistical significance of geographical clustering by comparing the mean distance to the means for up to 10,000,000 sets of randomly drawn non-carrier individuals matched by cohort status and number of parents with birthplace information available.

To assess whether PTVs or missense variants may be more geographically clustered than synonymous variants, we first identified a set of near-independent variants (r^2^>0.02) with MAC≥3 and MAF≤5% among the 14,874 unrelated individuals. This set included 4,312 PTVs, 91,851 missense variants, and 49,842 synonymous variants. For each variant, we computed the mean pairwise geographical distance in kilometers between the birthplaces across all pairs of the available parents of carriers of the minor allele and regressed this mean distance on variant class (PTVs, missense, or synonymous) and MAC, MAC^2^, and MAC^3^ (**Supplementary Table 14**).

We also assessed whether variants showing stronger enrichment (compared to NFE) are more likely to be geographically clustered. Starting with the three functional classes of variants identified above, we further restricted analysis to those variants found in gnomAD so we could calculate the enrichment in frequency over gnomAD NFE. We included 1,540 PTVs, 46,953 missense, and 28,912 synonymous variants in this analysis after pruning variants for LD with PLINK. As above, we computed the mean pairwise distances among parents of carriers of the minor allele and regressed mean distance on the logarithm of enrichment and MAC, MAC^2^, and MAC^3^ (**Supplementary Table 17**). In both analyses, we first assessed a model with the interaction terms but reported only the model without interactions if the interactions were not significant.

#### Heritability estimates and genetic correlations

We used genome-wide array genotype data on the 13,326 unrelated individuals for whom both exome sequence and array data were available to estimate heritability and genetic correlations for the 64 traits. We constructed a GRM with PLINK^105^ (v.1.90b, https://www.cog-genomics.org/plink2) by applying additional filters for MAF>1% and genotype missingness rate <2% to the set of previously-used genotyped SNVs, leaving 205,149 SNVs for GRM calculation. We used the exact mixed model approach of biMM^106^ (v.1.0.0, http://www.helsinki.fi/~mjxpirin/download.html) to estimate the heritability of our 64 traits and the genetic correlation of the 2,016 trait pairs.

## Figure Legends (Extended Data Figures)

**Extended Data Fig. 1. Comparison of allele frequencies of variants in FinMetSeq and NFE from gnomAD.** The comparison of allele frequencies shows the excess of variants at higher frequency in Finland as a result of the multiple bottlenecks experienced in Finnish population history.

**Extended Data Fig. 2. Proportional site frequency spectra between FinMetSeq and gnomAD NFE by variant annotation class.** In general, we find a depletion of the variants in the rarest frequency class, as well as enrichment of variants in the intermediate to common frequency range. The site frequency spectra were downsampled to 18,000 chromosomes for each dataset.

**Extended Data Fig. 3. Comparison of MAFs for trait-associated variants in FinMetSeq and NFE gnomAD.** Plotted in gray background is a 2-D histogram of variants with non-zero allele frequencies in both gnomAD and FinMetSeq but no trait associations. Variants significantly associated with at least one trait are colored and scaled proportionately to the association p-value, with more significant associations having a larger symbol. Variants >10X enriched in FinMetSeq compared to NFE are pink, those <10X enriched are in blue. The dashed line is the line of equal frequency. Variants unique to Finns and absent in gnomAD are not plotted.

**Extended Data Fig. 4. Gene-based association of extremely rare variants in *APOB* with serum total cholesterol.** The upper panel shows the distribution of the covariate adjusted and inverse-normal transformed phenotype. The lower panel displays the association statistics for each variant included in the gene-based test along with the trait value for minor allele carriers of each variant (orange triangles). SV.P is the P-value from the analysis of each variant in a single-variant analysis.

**Extended Data Fig. 5. Gene-based association of rare variants in *SECTM1* with HDL2 cholesterol.** The upper panel shows the distribution of the covariate adjusted and inverse-normal transformed phenotype. The lower panel displays the association statistics for each variant included in the gene-based test, along with the trait value for minor allele carriers of each variant (orange triangles). SV.P is the P-value from the analysis of each variant in a single-variant analysis.

**Extended Data Fig. 6. Gene-based association of extremely rare variants in *ABCA1* with serum HDL cholesterol.** The upper panel shows the distribution of the covariate adjusted and inverse-normal transformed phenotype. The lower panel displays the association statistics for each variant included in the gene-based test, along with the trait value for minor allele carriers of each variant (orange triangles). SV.P is the P-value from the analysis of each variant in a single-variant analysis.

**Extended Data Fig. 7. Gene-based association of extremely rare variants in *ALDH1L1* with glycine levels.** The upper panel shows the distribution of the covariate adjusted and inverse-normal transformed phenotype. The lower panel displays the association statistics for each variant included in the gene-based test, along with the trait value for minor allele carriers of each variant (orange triangles). SV.P is the P-value from the analysis of each variant in a single-variant analysis.

**Extended Data Fig. 8. Population structure of the FinMetSeq dataset, by region.** Population structure, by region, from principal components analysis of exome sequencing variant data (MAF > 1%), for 14,874 unrelated individuals whose parental birthplaces were known. Color indicates individuals with both parents born in the same region; gray indicates individuals with different parental birth regions, or missing information for one parent. Abbreviations for the regions: Usm, Uusimaa; Swf, Southwest Finland; Stk, Satakunta; Khm, Kanta-Hame; Prk, Pirkanmaa; Phm, Paijat-Hame; Kyl, Kymenlaakso; SKa, Southern Karelia; Nka, Northern Karelia; SSv, Southern Savonia; NSv, Northern Savonia; Ctf, Central Finland; SOs, Southern Ostrobothnia; Osb, Ostrobothnia; COs, Central Ostrobothnia; NOs, Northern Ostrobothnia; Kai, Kainuu; Lap, Lapland; x, split parental birthplaces. Large solid circles represent the center of each region. A map of Finland with regions labeled is supplied for reference.

**Extended Data Fig. 9. Hierarchical clustering tree produced by fineSTRUCTURE.** We identified 16 subpopulations within the FinMetSeq dataset by applying a haplotype-based clustering algorithm, fineSTRUCTURE, on 2,644 unrelated individuals born by 1955 whose parents were both born in the same municipality (Methods). Each subpopulation is named based on the most common parental birth location among its members, with the following abbreviations: NKa, North Karelia; NSv, North Savonia; SOs, South Ostrobothnia; NOs, North Ostrobothnia; Kai, Kainuu; Lap, Lapland; SuK, Surrendered Karelia. A map of Finland with regions labeled is supplied for reference. If multiple subpopulations share the same location label, the subpopulation is further distinguished with a numeral. NSv3 is used as an internal reference in enrichment analysis. See **Supplementary Table 15** for more detailed demographic descriptions of each subpopulation.

**Extended Data Fig. 10. Geographical clustering of missense and synonymous variants as a function of minor allele count and frequency enrichment over gnomAD NFE.** This represents the same analysis as Figure 5A, but for missense and synonymous variants rather than PTVs. Similar to PTVs, missense and synonymous variants that show greater enrichment in FinMetSeq are more likely to be geographically clustered. Blue and pink colors denote the frequency is lower or higher in FinMetSeq than in gnomAD NFE, respectively. The size of the point is proportional to the logarithm of the frequency ratio difference.

