## Extended Data Figures for "Exome sequencing identifies high-impact trait-associated alleles enriched in Finns"

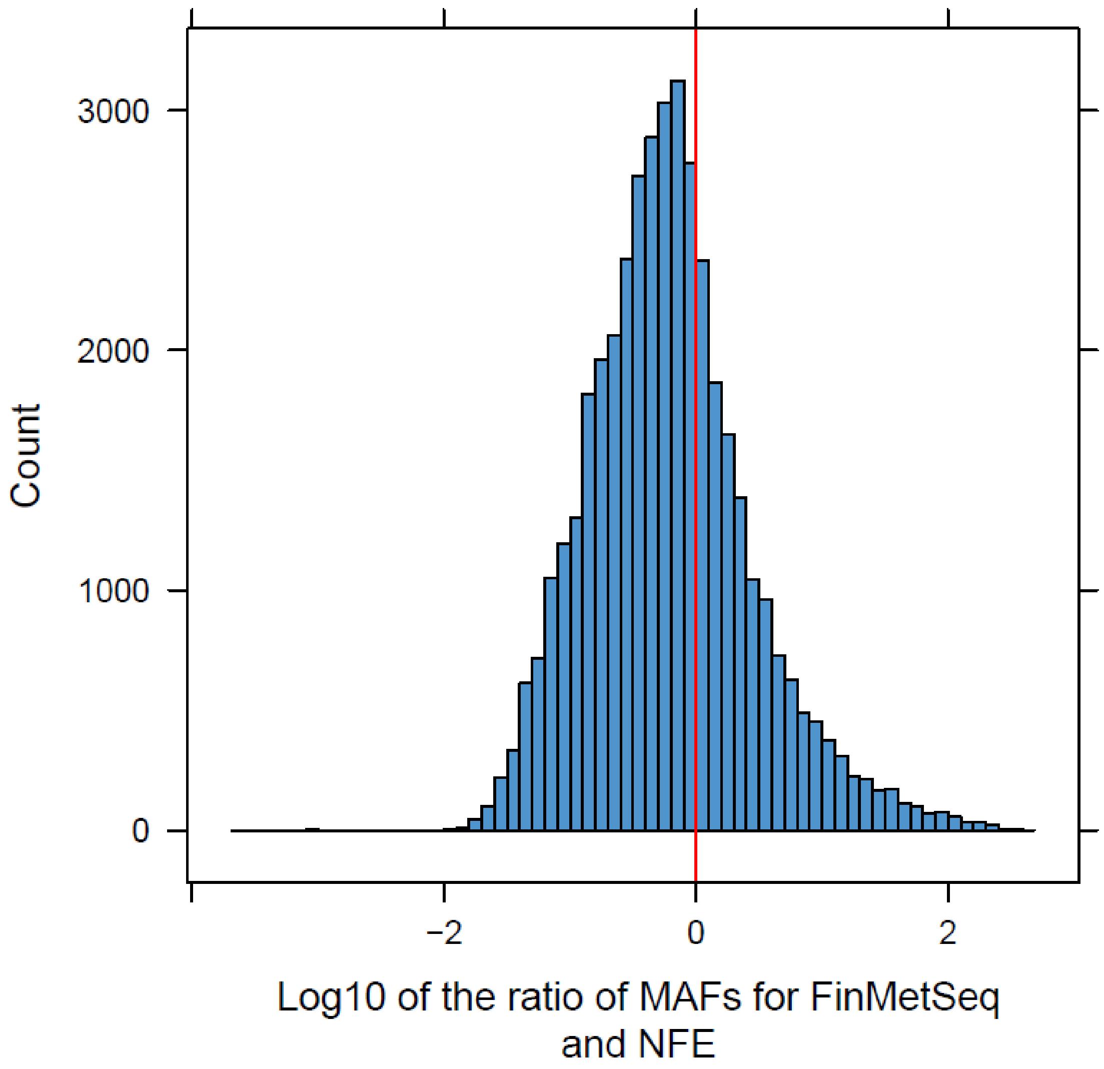


Extended Data Fig. 1. Comparison of allele frequencies of variants in FinMetSeq and NFE from gnomAD. The comparison of allele frequencies shows the excess of variants at higher frequency in Finland as a result of the multiple bottlenecks experienced in Finnish population history.

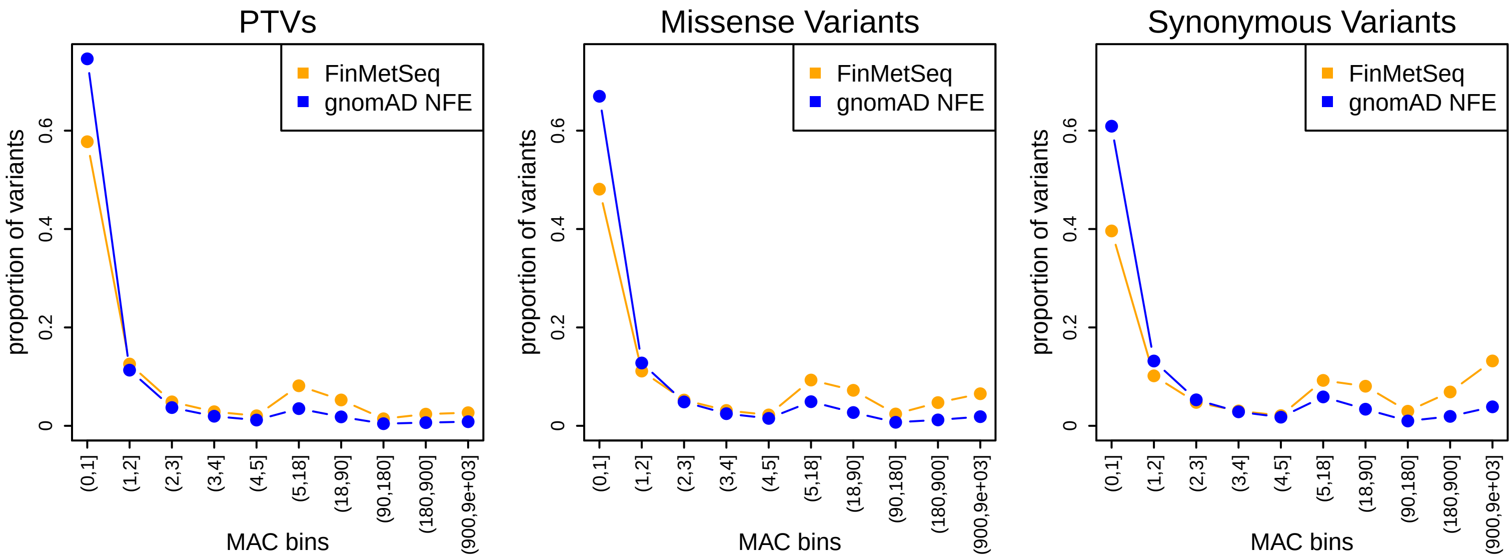


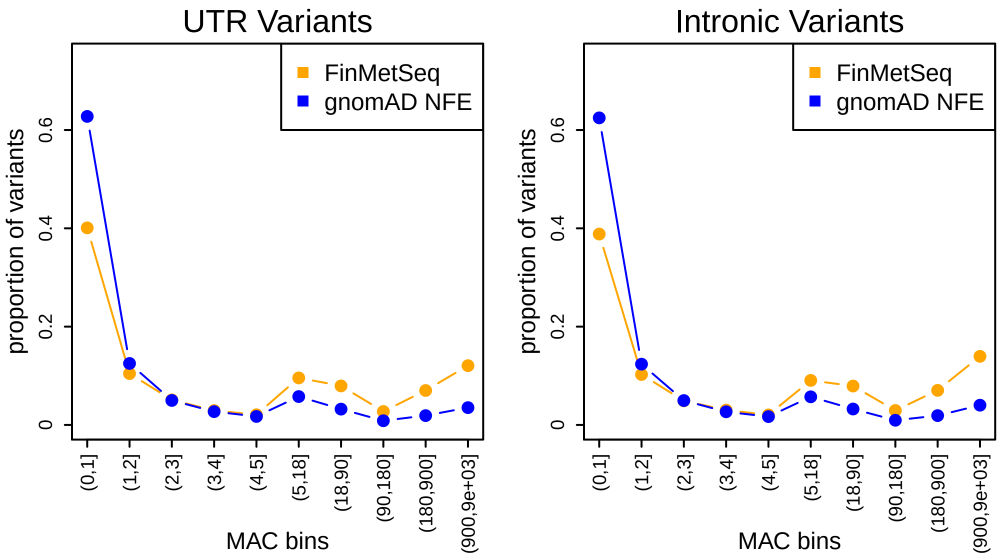


Extended Data Fig. 2. Proportional site frequency spectra between FinMetSeq and gnomAD NFE by variant annotation class. In general, we find a depletion of the variants in the rarest frequency class, as well as enrichment of variants in the intermediate to common frequency range. The site frequency spectra were down-sampled to 18,000 chromosomes for each dataset.

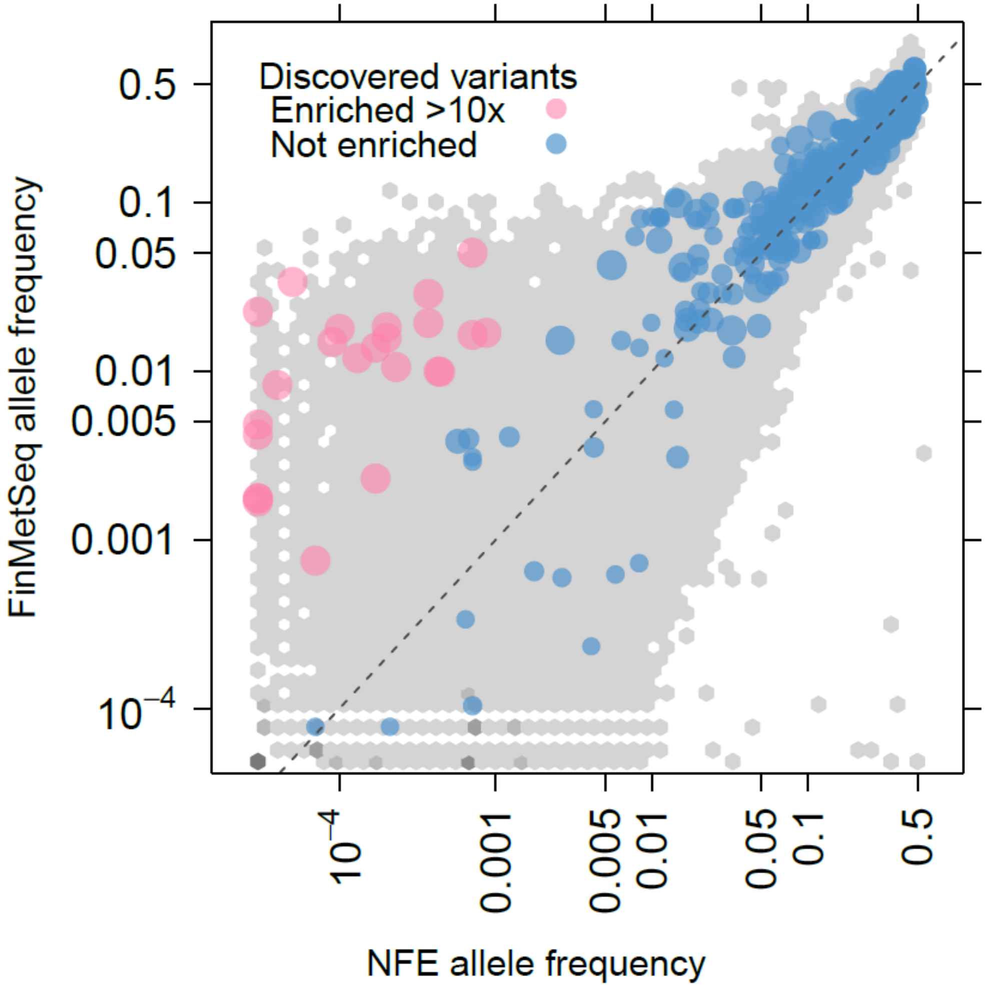


Extended Data Fig. 3. Comparison of MAFs for trait-associated variants in FinMetSeq and NFE in gnomAD. Plotted in gray background is a 2-D histogram of variants with non-zero allele frequencies in both gnomAD and FinMetSeq but no trait associations. Variants significantly associated with at least one trait are colored and scaled proportionately to the association p-value, with more significant associations having a larger symbol. Variants >10X enriched in FinMetSeq compared to NFE are pink, those <10X enriched are in blue. The dashed line is the line of equal frequency. Variants unique to Finns and absent in gnomAD are not plotted.

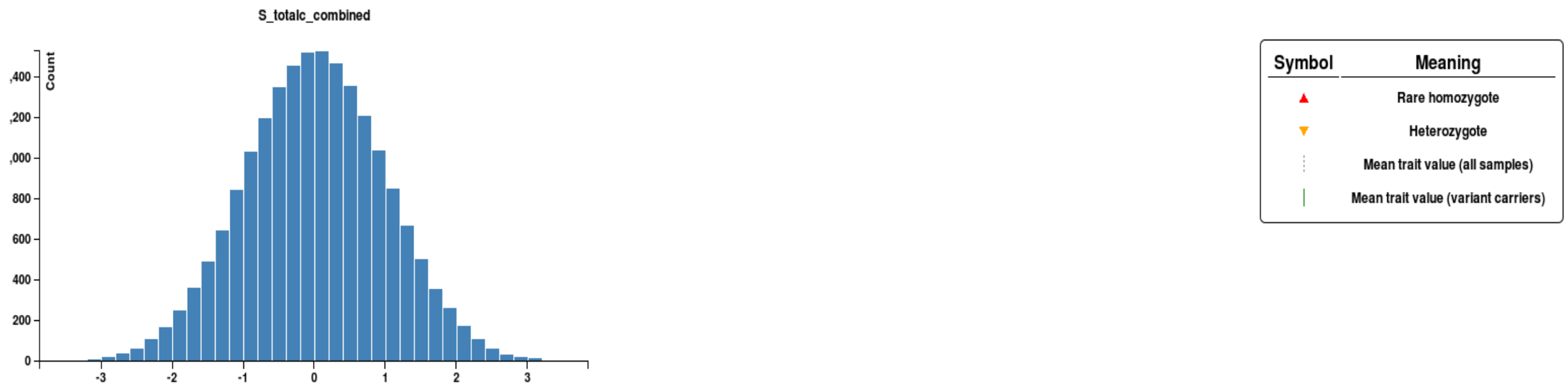


| GENE | mmskat.P | VARIANT | SV.P | BETA | MAF | MAC |
|---|---|---|---|---|---|---|
| APOB | 2.0e-13 | 2:21228642_G/A | 0.93 | 0.089 | 0.000030 | 1.0 |
| | | 2:21230094_AT/A | 0.0000013 | -2.8 | 0.000080 | 3.0 |
| | | 2:21230336_AT/A | 0.33 | -0.96 | 0.000030 | 1.0 |
| | | 2:21230583_C/A | 0.00011 | -2.7 | 0.000050 | 2.0 |
| | | 2:21231852_GA/G | 0.0065 | -2.7 | 0.000030 | 1.0 |
| | | 2:21233909_TGA/T | 0.16 | -1.4 | 0.000030 | 1.0 |
| | | 2:21234140_C/CG | 0.052 | -1.9 | 0.000030 | 1.0 |
| | | 2:21234858_C/CG | 0.0098 | -2.5 | 0.000030 | 1.0 |
| | | 2:21251412_T/C | 0.23 | -1.2 | 0.000030 | 1.0 |

Extended Data Fig 4. Gene-based association of extremely rare variants in APOB with serum total cholesterol. The upper panel shows the distribution of the covariate adjusted and inverse-normal transformed phenotype. The lower panel displays the association statistics for each variant included in the gene-based test along with the trait value for minor allele carriers of each variant (orange triangles). SV.P is the P-value from the analysis of each variant in a single-variant analysis.

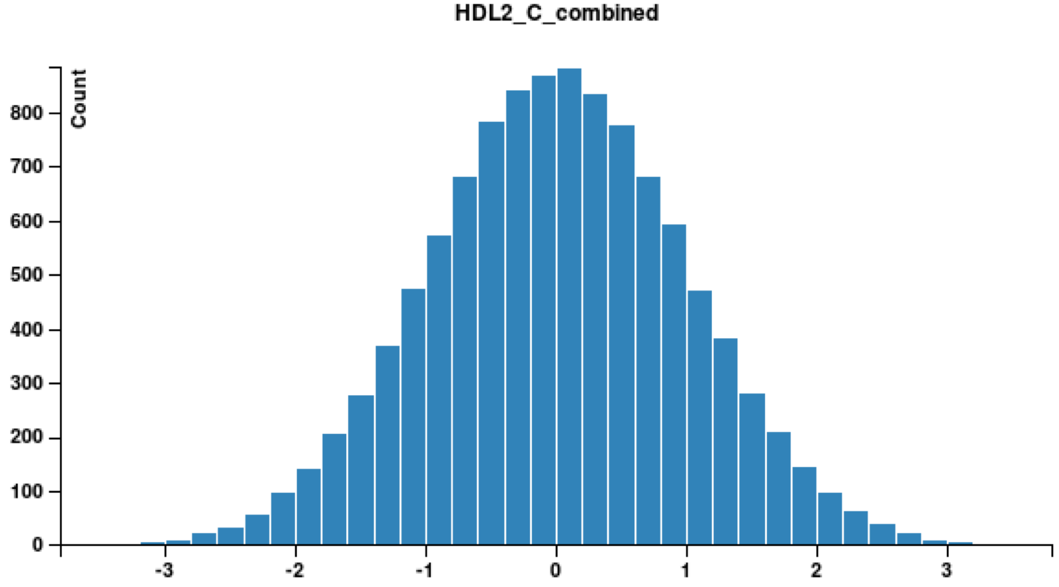


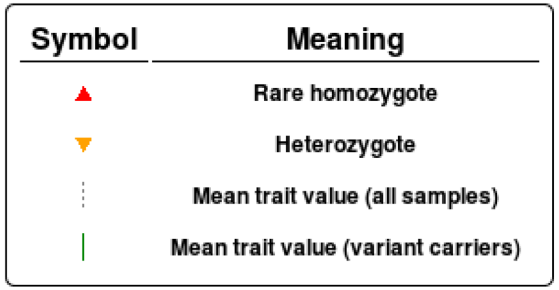


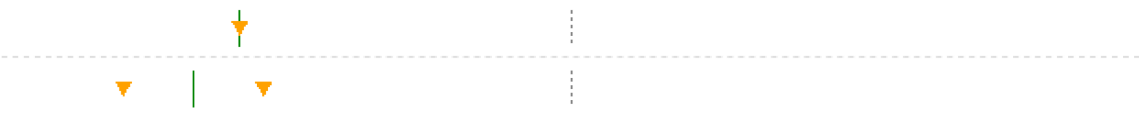

| GENE | mmskat.P | VARIANT | SV.P | BETA | MAF | MAC |
|---|---|---|---|---|---|---|
| SECTM1 | 5.6e-7 | 17:80280768_G/A | 0.0088 | -2.6 | 0.000050 | 1.0 |
| | | 17:80280876_G/A | 0.000020 | -2.9 | 0.000090 | 2.0 |

Extended Data Fig. 5. Gene-based association of rare variants in SECTM1 with HDL2 cholesterol. The upper panel shows the distribution of the covariate adjusted and inverse-normal transformed phenotype. The lower panel displays the association statistics for each variant included in the gene-based test, along with the trait value for minor allele carriers of each variant (orange triangles). SV.P is the P-value from the analysis of each variant in a single-variant analysis.

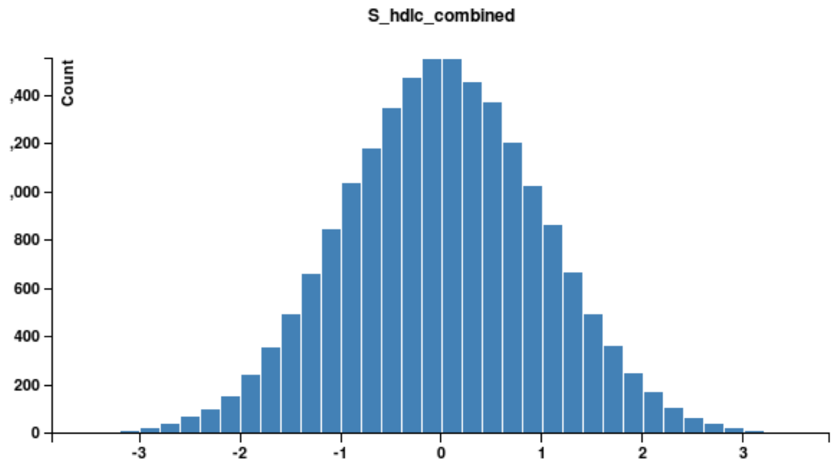


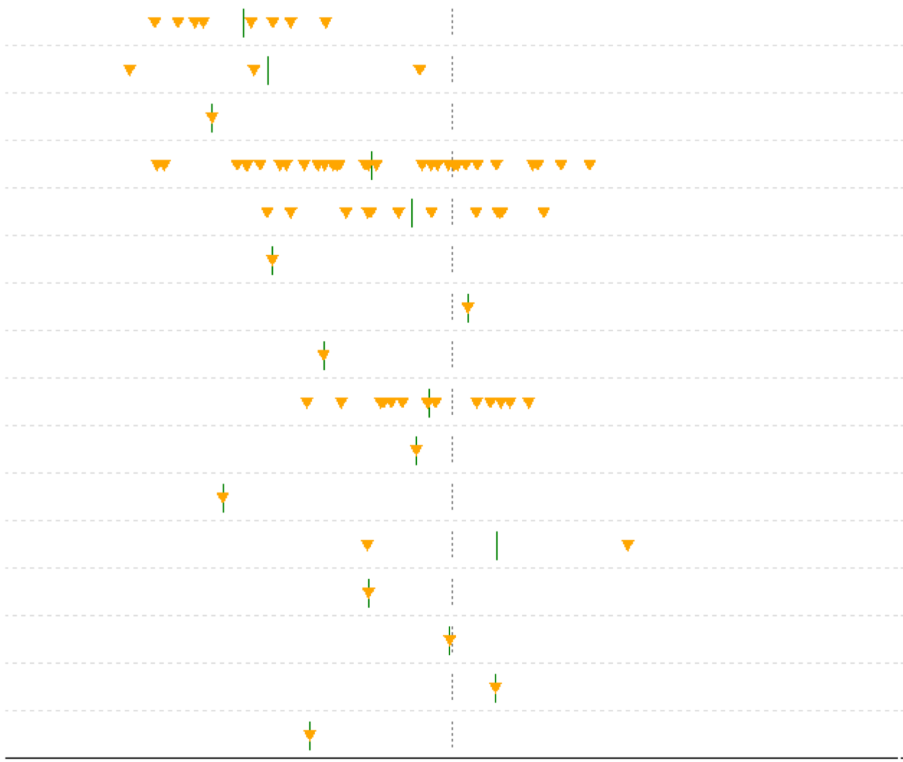

| GENE | mmskat.P | VARIANT | SV.P | BETA | MAF | MAC |
|---|---|---|---|---|---|---|
| ABCA1 | 5.2e-13 | 9:107548661_A/G | 4.8e-10 | -2.0 | 0.00023 | 9.0 |
| | | 9:107550254_G/A | 0.00020 | -2.1 | 0.000080 | 3.0 |
| | | 9:107550798_T/G | 0.035 | -2.1 | 0.000030 | 1.0 |
| | | 9:107558416_T/C | 4.3e-7 | -0.88 | 0.00083 | 32 |
| | | 9:107560784_C/T | 0.26 | -0.31 | 0.00034 | 13 |
| | | 9:107568659_G/A | 0.054 | -1.9 | 0.000030 | 1.0 |
| | | 9:107578515_C/A | 0.62 | 0.48 | 0.000030 | 1.0 |
| | | 9:107586839_G/A | 0.24 | -1.1 | 0.000030 | 1.0 |
| | | 9:107589238_C/G | 0.53 | -0.16 | 0.00036 | 14 |
| | | 9:107591298_G/A | 0.89 | 0.13 | 0.000030 | 1.0 |
| | | 9:107593339_G/A | 0.032 | -2.1 | 0.000030 | 1.0 |
| | | 9:107593907_C/A | 0.57 | 0.39 | 0.000050 | 2.0 |
| | | 9:107593948_C/T | 0.40 | -0.82 | 0.000030 | 1.0 |
| | | 9:107599789_A/G | 0.83 | 0.21 | 0.000030 | 1.0 |
| | | 9:107646739_G/A | 0.66 | 0.42 | 0.000030 | 1.0 |
| | | 9:107646741_GC/G | 0.27 | -1.1 | 0.000030 | 1.0 |

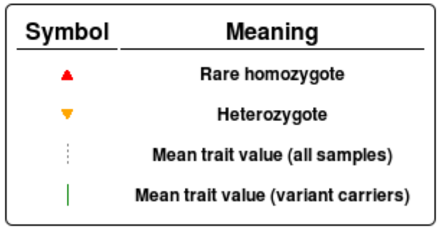


Extended Data Fig 6. Gene-based association of extremely rare variants in ABCA1 with serum HDL cholesterol. The upper panel shows the distribution of the covariate adjusted and inverse-normal transformed phenotype. The lower panel displays the association statistics for each variant included in the gene-based test, along with the trait value for minor allele carriers of each variant (orange triangles). SV.P is the P-value from the analysis of each variant in a single-variant analysis.

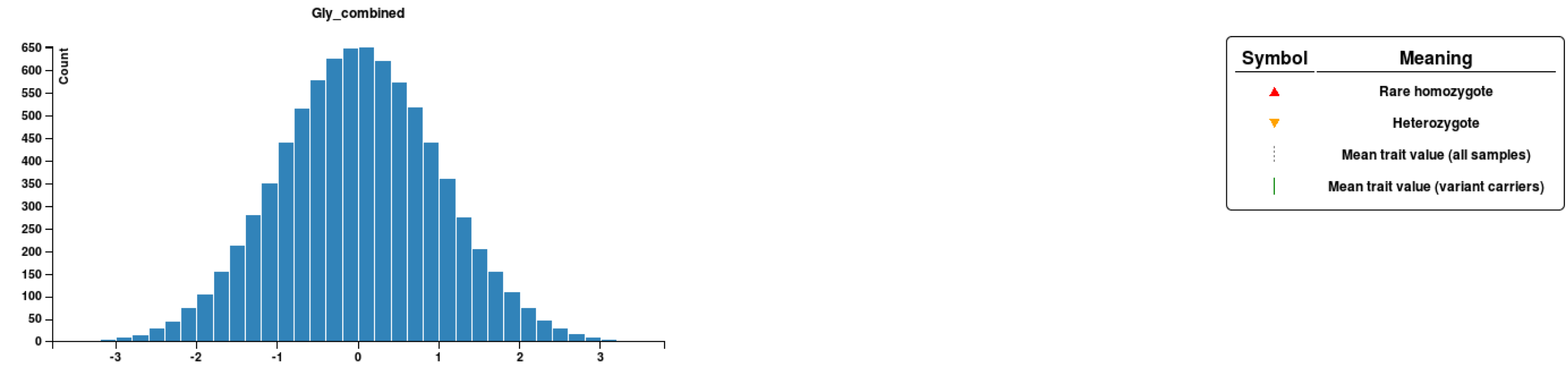


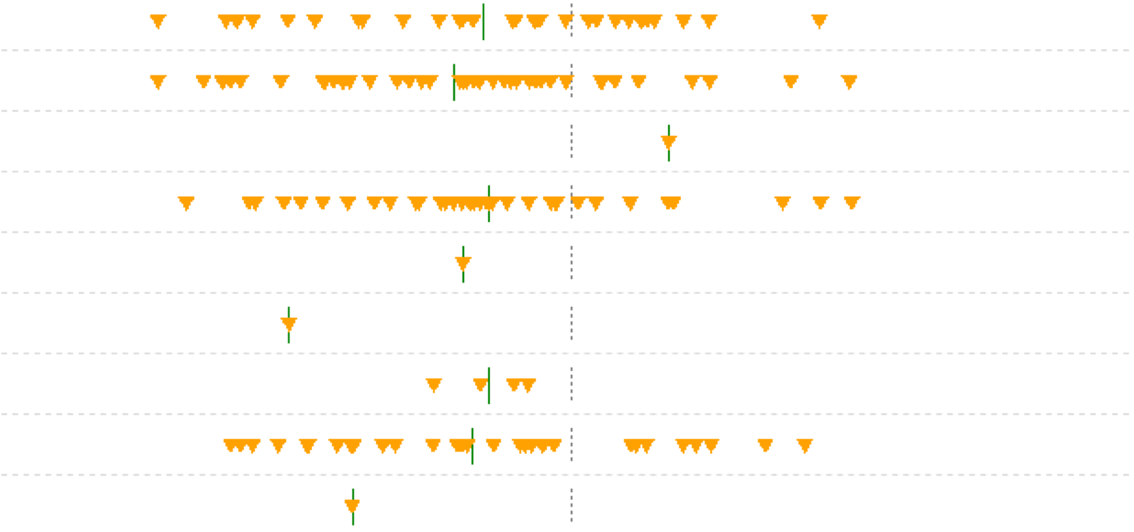

| GENE | mmskat.P | VARIANT | SV.P | BETA | MAF | MAC |
|---|---|---|---|---|---|---|
| ALDH1L1 | 3.9e-21 | 3:125828918_G/A | 0.000029 | -0.73 | 0.0020 | 33 |
| | | 3:125831672_G/A | 1.8e-8 | -0.87 | 0.0026 | 42 |
| | | 3:125850306_T/C | 0.53 | 0.62 | 0.000060 | 1.0 |
| | | 3:125854377_C/T | 0.000072 | -0.64 | 0.0024 | 39 |
| | | 3:125873443_C/T | 0.44 | -0.76 | 0.000060 | 1.0 |
| | | 3:125877290_G/A | 0.039 | -2.1 | 0.000060 | 1.0 |
| | | 3:125879703_G/T | 0.19 | -0.66 | 0.00024 | 4.0 |
| | | 3:125879755_C/T | 0.000050 | -0.70 | 0.0021 | 34 |
| | | 3:125879807_T/A | 0.13 | -1.5 | 0.000060 | 1.0 |

Extended Data Fig. 7. Gene-based association of extremely rare variants in ALDH1L1 with glycine levels. The upper panel shows the distribution of the covariate adjusted and inverse-normal transformed phenotype. The lower panel displays the association statistics for each variant included in the gene-based test, along with the trait value for minor allele carriers of each variant (orange triangles). SV.P is the P-value from the analysis of each variant in a single-variant analysis.

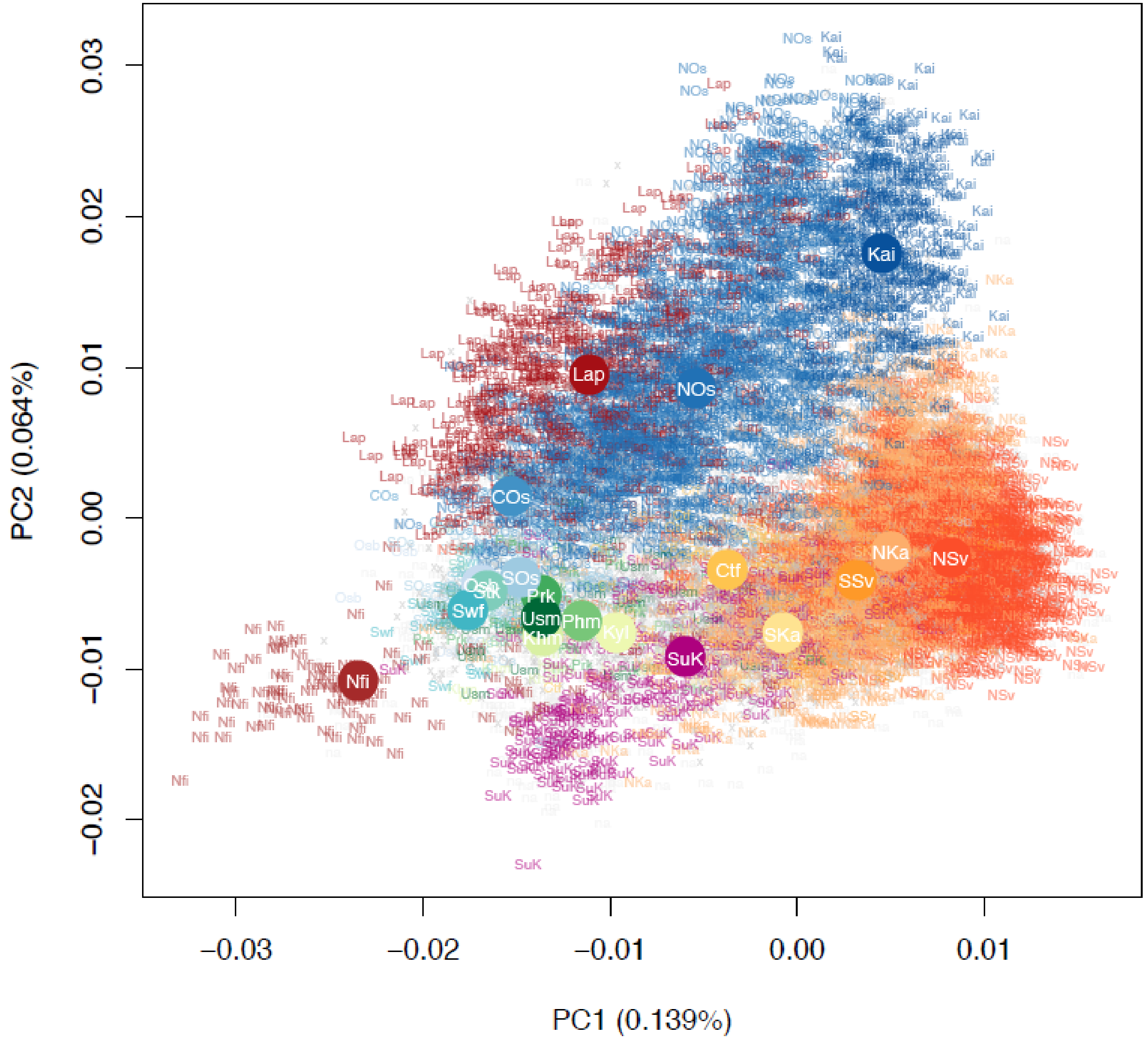


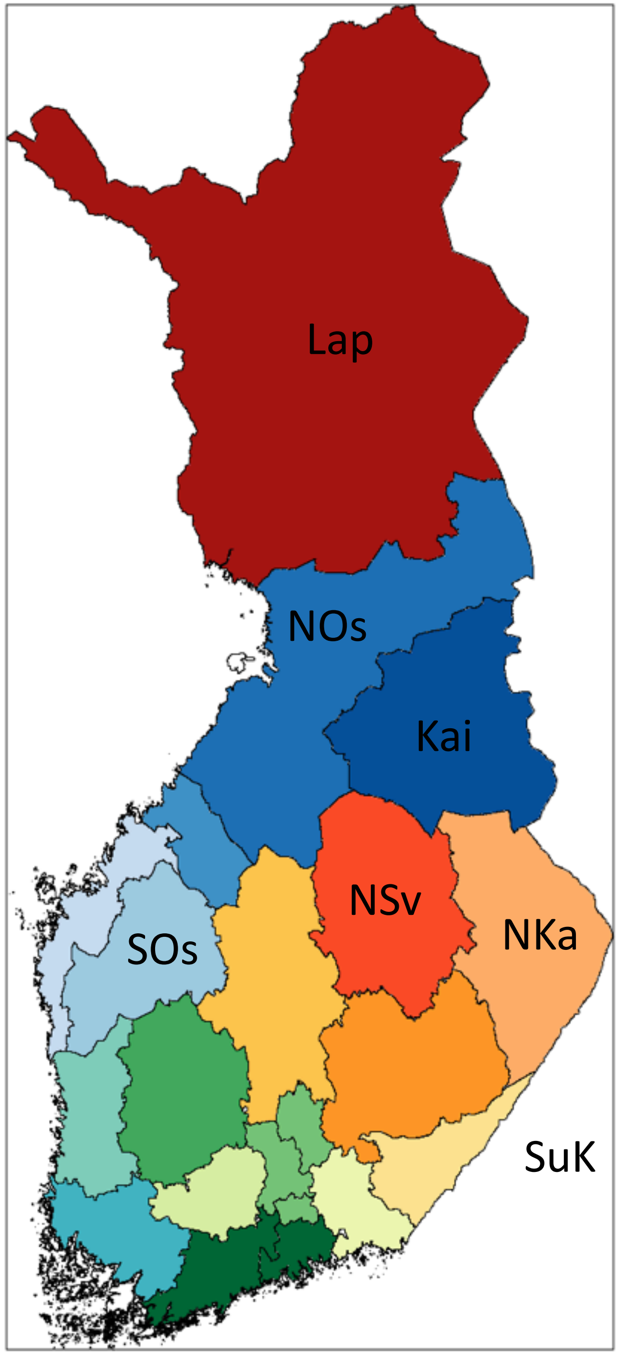


Extended Data Figure 8. Population structure of the FinMetSeq dataset, by region. Population structure, by region, from principal components analysis of exome sequencing variant data (MAF > 1%), for 14,874 unrelated individuals whose parental birthplaces were known. Color indicates individuals with both parents born in the same region; gray indicates individuals with different parental birth regions, or missing information for one parent. Abbreviations for the regions: Usm, Uusimaa; Swf, Southwest Finland; Stk, Satakunta; Khm, Kanta-Hame; Prk, Pirkanmaa; Phm, Paijat-Hame; Kyl, Kymenlaakso; SKa, Southern Karelia; Nka, Northern Karelia; SSv, Southern Savonia; NSv, Northern Savonia; Ctf, Central Finland; SOs, Southern Ostrobothnia; Osb, Ostrobothnia; COs, Central Ostrobothnia; NOs, Northern Ostrobothnia; Kai, Kainuu; Lap, Lapland; x, split parental birthplaces. Large solid circles represent the center of each region. A map of Finland with regions labeled is supplied for reference.

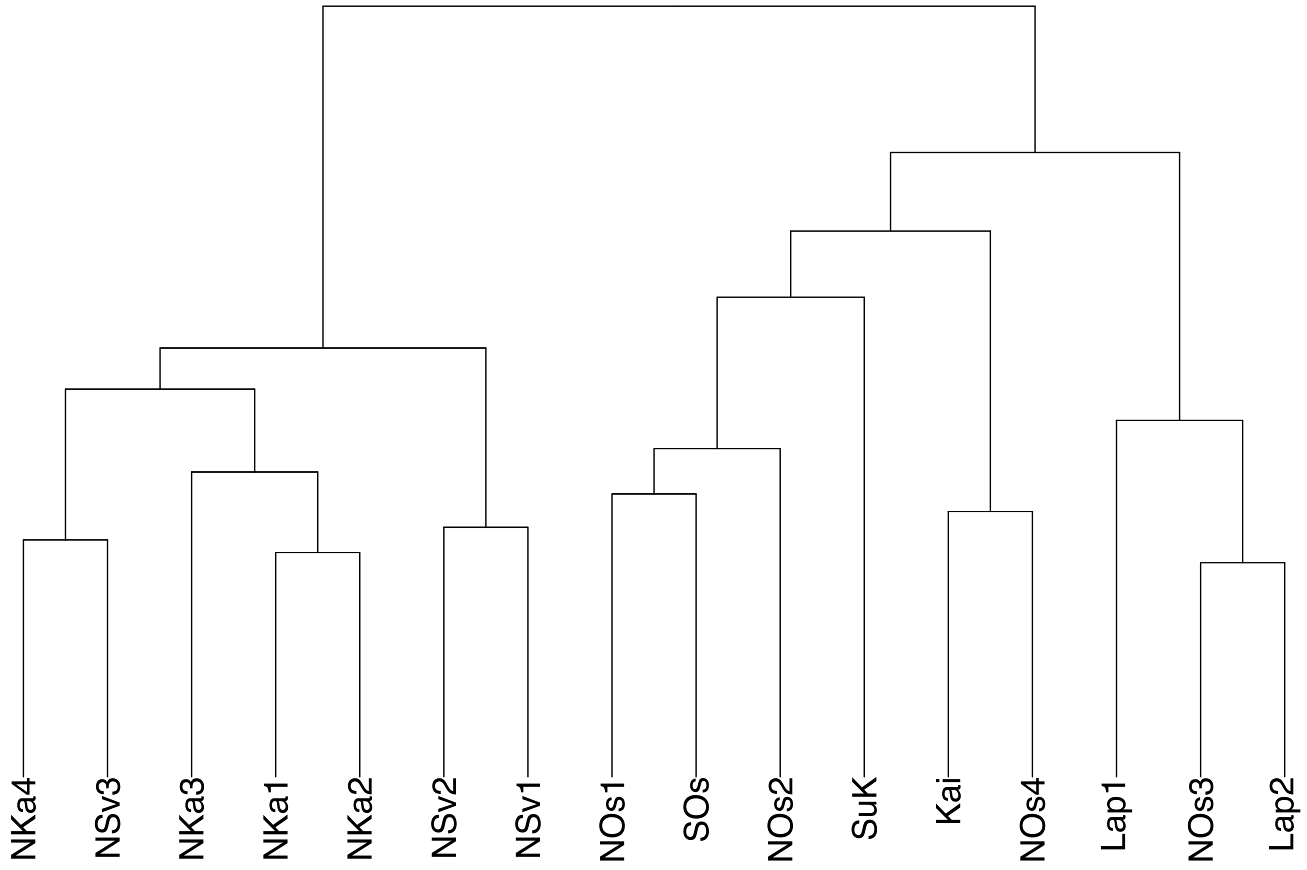


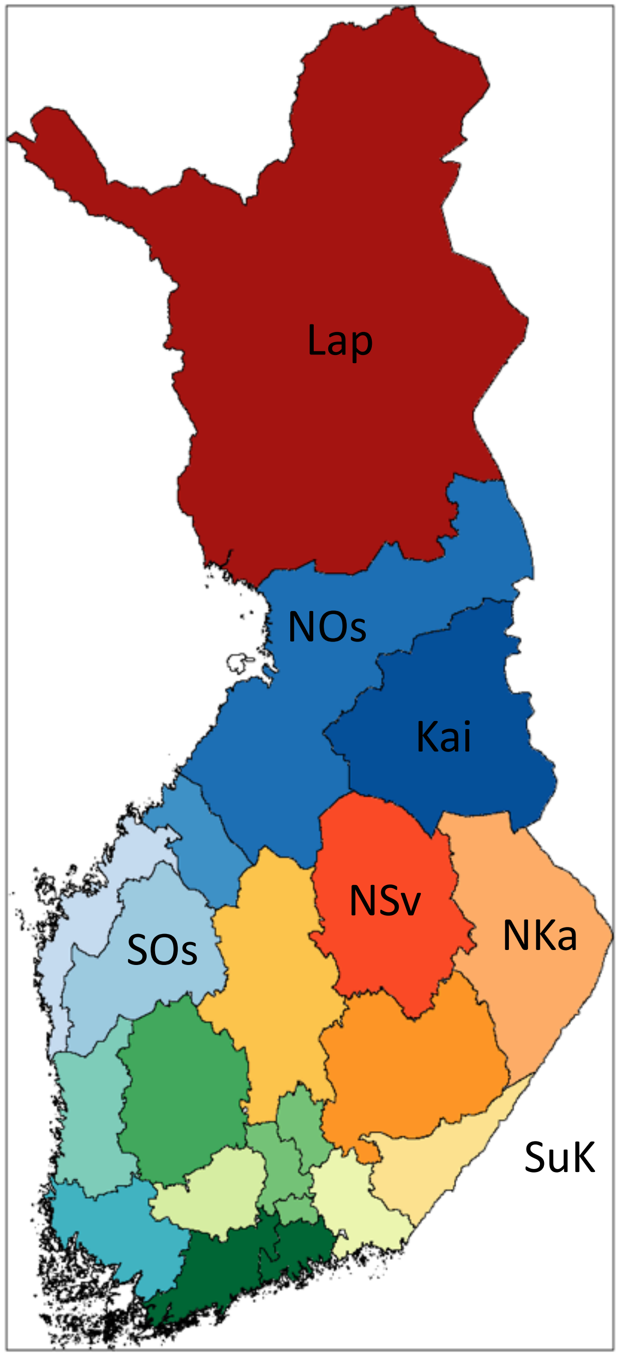


Extended Data Fig. 9. Hierarchical clustering tree produced by fineSTRUCTURE. We identified 16 subpopulations within the FinMetSeq dataset by applying a haplotype-based clustering algorithm, fineSTRUCTURE, on 2,644 unrelated individuals born by 1955 whose parents were both born in the same municipality (Methods). Each subpopulation is named based on the most common parental birth location among its members, with the following abbreviations: NKa, North Karelia; NSv, North Savonia; SOs, South Ostrobothnia; NOs, North Ostrobothnia; Kai, Kainuu; Lap, Lapland; SuK, Surrendered Karelia. A map of Finland with regions labeled is supplied for reference. If multiple subpopulations share the same location label, the subpopulation is further distinguished with a numeral. NSv3 is used as an internal reference in enrichment analysis. See Supplementary Table 15 for more detailed demographic descriptions of each subpopulation.

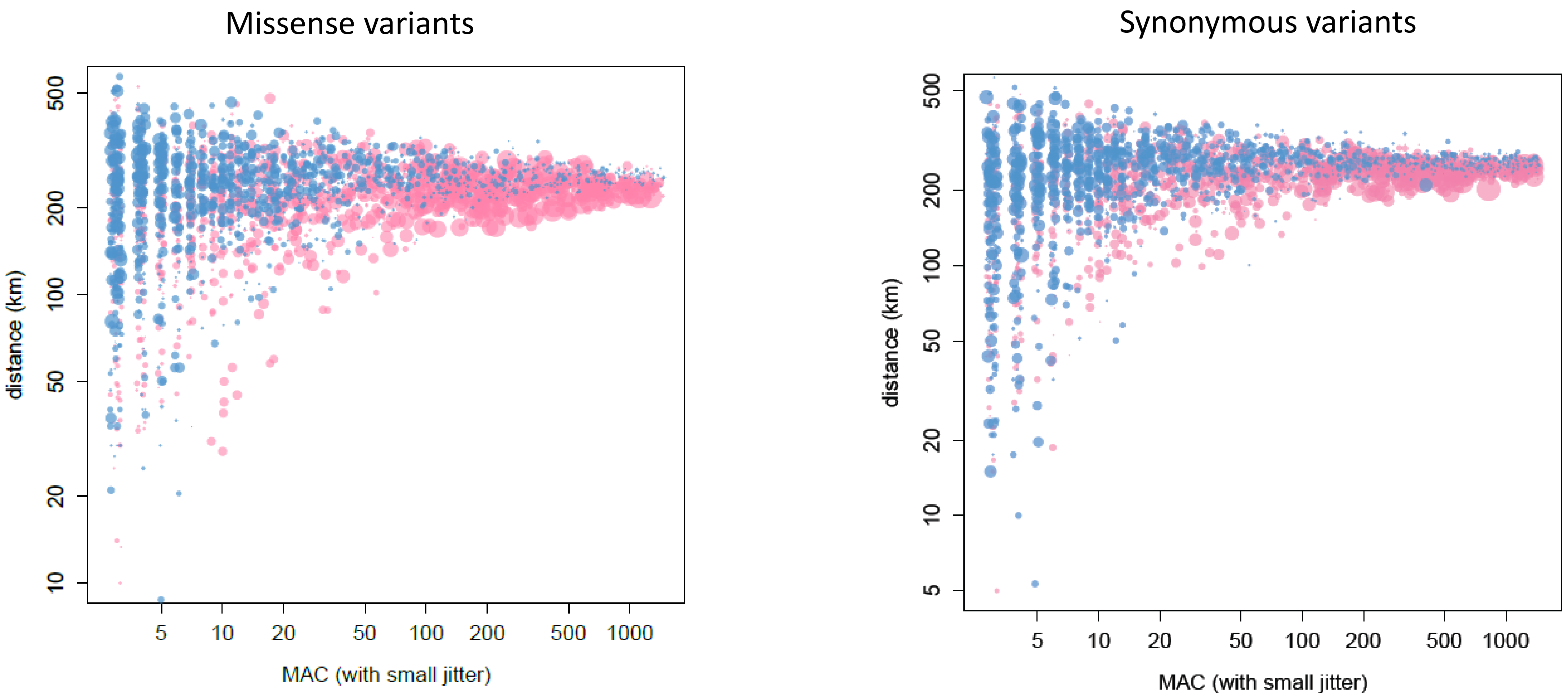


Extended Data Fig. 10. Geographical clustering of missense and synonymous variants as a function of minor allele count and frequency enrichment over gnomAD NFE. This represents the same analysis as Figure 5A, but for missense and synonymous variants rather than PTVs. Similar to PTVs, missense and synonymous variants that show greater enrichment in FinMetSeq are more likely to be geographically clustered. Blue and pink colors denote the frequency is lower or higher in FinMetSeq than in gnomAD NFE, respectively. The size of the point is proportional to the logarithm of the frequency ratio difference.
