## Supplementary Information for "Exome sequencing identifies high-impact trait-associated alleles enriched in Finns"

Supplementary Results: Novel association descriptions	Page 2-5

Supplementary Methods: Multiplicity adjustment	Page 6-9

Supplementary Methods: Power calculation	Page 10

Supplementary References	Page 11

**Supplementary Results**

**Novel variants highly enriched in Finland**

*Anthropomorphic traits*

Two very rare variants, rs757565850 (p.Lys473Asn), a predicted deleterious missense variant in *BOD1L1* (P=4.7×10^-7^, $\hat{\beta}$=-2.50), and 5:140181423:CA/C, a frameshift variant in *PCDHA3* (P=2.7×10^-7^, $\hat{\beta}$=2.56), have very strong effects on waist-hip ratio (WHR, **Table 2**). *BOD1L1* helps regulate genome integrity as a component of a replication fork protection pathway^1^, while *PCDHA3* is a member of the protocadherin alpha gene cluster thought to regulate neuronal migration during cortical development^2^. rs757565850 looks to be unique to Finns, while the *PCDHA3* deletion is not in gnomAD. WHR is increasingly considered more sensitive than BMI as a predictor of cardiovascular disease and diabetes risk^3,4^. It is not obvious why variation in either gene should have such a strong effect on WHR. We interpret these associations cautiously, given the absence of replication evidence or obvious biological significance.

*HDL-C*

A *LIPG* splice acceptor variant (rs200435657), present at 0.25% in FinMetSeq but not observed in NFE, is associated in the combined analysis with HDL2-C (P_meta_=5.6×10^-10^, $\hat{\beta}$=0.62), phosphatidylcholine (P_meta_=1.1×10^-8^, $\hat{\beta}$=0.58), and total phosphatidylglycerol (P_meta_=1.1×10^-7^, $\hat{\beta}$=0.54; **Table 2**). At *NR1H3*, we identified an independent association of a predicted deleterious missense variant (rs199947343, p.Met213Val) 140X more frequent in FinMetSeq than in NFE controls, with HDL-C in FinMetSeq (P=1.4×10^-7^, $\hat{\beta}$=0.43; **Table 2**), and with VLDL-C (P_meta_=3.1×10^-7^, $\hat{\beta}$=-0.41) and HDL2-C (P_meta_=1.3×10^-8^, $\hat{\beta}$=0.46) in the combined analysis (**Table 2**). At *CETP*, we identified a small deletion (rs751916721) disrupting a splice donor site that is 56.7X more frequent in FinMetSeq than in NFE associated with HDL-C (P=1.1×10^-14^, $\hat{\beta}$=0.95) and serum ApoA1 levels (P=2.6×10^-8^, $\hat{\beta}$=0.83), independent of previous associations at this locus (**Table 2**).

At *ANGPTL8* (also called C19orf80) we identified a stop gain variant (rs760351239, p.Gln131X) not seen in NFE, associated with HDL2-C (P_meta_=1.1×10^-8^, $\hat{\beta}$=0.57) in the combined analysis (**Table 2**). This appears to be independent of a previously reported variant associated with decreased HDL-C in Mexican-Americans.

At *DPEP3* we identified a predicted deleterious missense variant (rs200922436, p.Arg154Lys) in 22.5X enriched in FinMetSeq associated with decreased HDL-C (P=1.6×10^-7^, $\hat{\beta}$=-0.30; **Table 2**) and decreased ApoA1 (P_meta_=4.0×10^-7^, $\hat{\beta}$=-0.25; **Table 2**).

*Apolipoprotein B*

We identified a predicted deleterious missense variant (rs369295276, p.418Asn) in *AP1M2* that is 167X more frequent in FinMetSeq than in NFE associated with decreased serum ApoB levels (P=5.8×10^-8^, $\hat{\beta}$=-0.28) in FinMetSeq (**Table 2**), and with total cholesterol in intermediate density lipoprotein (IDL-C), concentration of IDL particles (IDL-P), and remnant cholesterol in the combined analysis (**Table 2**). *AP1M2* encodes a subunit of the heterotetrameric adaptor-related protein complex 1a that plays a role in protein sorting in the trans-Golgi network. This association is independent of previously reported association of this locus with thrombosis^5^.

*Glucose metabolism and related traits*

A predicted deleterious missense variant in *G6PC* (rs201961848, p.Ala204Ser) is present at 2.5% frequency in FinMetSeq but absent in NFE. *G6PC* encodes glucose-6-phosphatase, which catalyzes the production of free glucose from glycogen in the liver and other organs. Autosomal recessive transmission of any of ~100 previously reported *G6PC* mutations result in glycogen storage disease type 1a, characterized by short stature, hypertriglyceridemia, hyperuricemia, lactic acidosis, and hypoglycemia (https://www.omim.org/entry/232200). Eight individuals in FinMetSeq are homozygous for this novel variant. Heterozygous carriers of deleterious *G6PC* variants in the DiscovEHR cohort also exhibited hypertriglyceridemia, increased uric acid, and increased risk of gout^6^. We found significant associations in heterozygous carriers with several traits; in FinMetSeq, rs201961848 was associated with monounsaturated fatty acid (MUFA) levels (P=4.4 ×10^-7^, $\hat{\beta}$=0.27; **Table 2**), and in the combined analysis with glycerol (P_meta_=4.1×10^-7^, $\hat{\beta}$=0.18), total triglycerides (P_meta_=1.3×10^-7^, $\hat{\beta}$=0.20), and plasma C-reactive protein (CRP) (P_meta_=4.0×10^-9^, $\hat{\beta}$=0.19; **Table 2**).

**Novel variants not highly enriched in Finland**

Given the large sample size and number of traits assessed, we identified novel associations even to variants that had similar or only moderately increased frequencies in FinMetSeq compared to NFE. For example, a missense variant in *EPB41L5* (rs28930677, p.His334Tyr) that is twice as frequent in FinMetSeq as in NFE showed significant association to indices of renal function in the combined analysis (Creatinine: P_meta_=2.5×10^-12^, $\hat{\beta}$=0.098; eGFR: P_meta_=4.8×10^-12^, $\hat{\beta}$=-0.11; **Table 2**). This variant was suggestively (P=2.0×10^-6^) associated with kidney function in a recent GWAS of ~110,000 individuals, but at half the frequency (MAF=0.047) that we observed in FinMetSeq^7^. *EPB41L5* is highly expressed in the kidney and functions as a podocyte focal adhesion component required for podocyte stability at the kidney filtration barrier. Podocyte detachment from the glomerular basement membrane is characteristic of late stage glomerular disease. Deletion of its ortholog *Epb41l5* in mice led to proteinuria, detachment of podocytes, and development of focal segmental glomerulosclerosis^8^.

**Supplementary Methods: Multiplicity adjustments in an exome-wide screen for multiple phenotypes**

### Motivation and background

#### *The choice of False Discovery Rate (FDR) as a criterion*

The goal of studies as comprehensive as this one is to generate hypotheses regarding which genetic variation has impact on the phenotypes of interest, and to identify the associations that are real. This goal is formalized by the criterion of FDR control: procedures that control FDR assure that, on average, the proportion of false discoveries among the one highlighted is arbitrarily small. It is useful to contrast this with two other options: 1) family wise error rate control (FWER) and 2) no adjustment for multiplicity. The goal of FWER is to assure that the probability of making any false discovery is arbitrarily small. When studying a large enough number of heritable quantitative traits one expects many true discoveries, and it is therefore sensible to tolerate the inclusion of some false ones. On the other hand, to entirely ignore the multiplicity of phenotypes we analyze is equivalent to exposing ourselves (and our field) to the risk of many false discoveries.

#### *Strong exome-wide selection and large power across phenotypes*

The primary contribution to multiple testing in our study is variation across the exome (up to 603,057 variants tested). The multiplicity of traits is a much smaller contributor (64 traits). The strategy we selected to adjust for multiplicity is mindful of this fact and protects us more stringently from the most likely error, that of variants that have no effect on any phenotype, than from erroneously attributing to a truly trait-associated variant an influence on an additional trait.

#### *Adaptive versus universal threshold*

A characteristic of FDR controlling procedures is that they reject hypotheses with a p-value smaller then some critical value $p^{*}$ which depends not only on the target value for the FDR and number of hypotheses tested, but also on the number of association signals in the dataset. In other words, different collections of p-values referring to the same number of hypotheses are going to result in different thresholds $p^{*}$. This is a powerful feature of FDR controlling procedures, but one that might lead to some confusion in genetic association studies where we are used to “genome-wide” thresholds. To bridge this gap, we have selected for consideration as potentially meaningful association signals only the ones relative to variants whose smallest p-value for association across the 64 traits is smaller than 5×10^-7^.

#### *Prior work we leverage*

We rely on the results of two prior papers dealing with FDR control in genetic studies and in the presence of multiple phenotypes. Brzyski et al. illustrated how to achieve FDR control when working with a subset of hypotheses selected on the basis of the data, so that one can select the “most promising” hypotheses^9^ (as defined above). Peterson et al. described a powerful multi-phenotype FDR controlling procedure which takes a less conservative approach across phenotypes^10^.

### Multiplicity adjustment procedure

The procedure we adopt has two stages.

Stage 1: Identify the variants that are associated to at least one phenotype, controlling the FDR at level 0.05 (non-null variants).

Stage 2: Identify the phenotypes associated to the variants discovered in step 1, controlling the expectation of the average false discovery proportion across all such non-null variants.

In the interest of clarity, let $p_{ij}$ be the p-value for association between variant $i$ and trait $j$ (as obtained with EMMAX), where $i=1,\ldots, 603,057$ and $j=1,\ldots,64$ covering all variants tested for at least one trait and covering all traits.

#### *Stage 1: identify non-null variants*

To focus our attention on variants that are likely to be important, bridge the gap between FDR controlling procedures and fixed significance threshold methods, and mitigate the difficulties that linkage disequilibrium poses for FDR methods, we adopted a two-step procedure:

I. Screen all variants to identify a subset R of variants that have a p-value for association to at least one trait smaller than $\boldsymbol{5\times}\boldsymbol{10}^{\boldsymbol{-7}}$: $\text{R}\boldsymbol{=\{i:}\mathbf{min}_{\boldsymbol{j}}\boldsymbol{p}_{\boldsymbol{ij}}\boldsymbol{\leq5\times}\boldsymbol{10}^{\boldsymbol{-7}}\boldsymbol{\}}$.

II. Discover non-null variants at FDR<0.05. To achieve this objective we proceed as follows.

(a) For each variant in R calculate the Simes’ p-value for the hypothesis of no association to any trait: $p_{i}^{s}=\min_{j}p_{im}\times\frac{64}{m}$, where $p_{i1}\leq p_{i2}\leq\cdots p_{i64}$ is the set of ordered p-values for association between variant $i$ and each of the 64 traits. The Simes’ method combines p-values to test a global null that is valid also when the p-values are dependent; in our case the global null is the hypothesis that variant $i$ is not associated with any of the 64 traits.
(b) Apply the Benjamini Hochberg (BH) procedure^11^ to the p-values $\{p_{i}^{s},i\in\text{R}\}$ at a level $0.05\times|\text{R}|/603,057$ to identify the set S of non-null variants with FDR $\leq0.05$. Note that the BH procedure has been applied at a level more stringent than $0.05$ to correct for the fact that the set of variants R was selected on the basis of the data. The validity of this procedure is described in Brzyski et al.^9^.

#### *Stage 2: discovery of associated phenotypes*

To determine which traits are associated to the non-null variants in S we apply the Benjamini and Bogomolov procedure^12^, whose efficacy for multi-trait genetic association studies was explored in Peterson et al.^10^. Specifically, for each $i\in\text{S}$, we apply the BH procedure to the set of p-values $\{p_{i1},\ldots,p_{i64}\}$ at a level $0.05|\text{S}|/603,057$, which guarantees that the expected value of the average false discovery proportion across all $i\in\text{S}$ is smaller than $0.05$. Analyzing the associations variant by variant (as opposed to pooling all the p-values together) results in an increase of power to detect the association of variants with effects on multiple phenotypes^10^.

**Supplementary Methods: Power calculation for Figures 3A & 3B**

Let *n* be the number of subjects in an association test of a single variant with a single trait. Suppose that *s* subjects have exactly one copy of the minor allele and the remaining *n-s* have none. Let *y_i_* and *x_i_* denote the phenotype and genotype of subject *i* for this trait and variant. The *y_i_* are standardized to have mean zero and variance one and the *x_i_* are coded as minor allele count shifted by the mean $\frac{s}{n}=2\hat{q}$, where $\hat{q}$ is the minor allele frequency (MAF). The ordinary least squares (OLS) regression model is then, in vector notation, **y** = *β***x** + **ε**, where the entries of **ε** are independent and identically distributed N(0,$\sigma^{2}$).

After obtaining the OLS estimates $\hat{\beta}$ of *β* and $\hat{\sigma}$ of *σ*, we compute the statistic $t=\hat{\beta}/\mathrm{se}(\hat{\beta})$, where $\mathrm{se}\left( \hat{\beta} \right)=\hat{\sigma}/\sqrt{\mathbf{x}^{\boldsymbol{T}}\mathbf{x}}$. Because **x** has been centered, $\mathbf{x}^{\boldsymbol{T}}\mathbf{x}=s{(1-2\hat{q})}^{2}+(n-s){(2\hat{q})}^{2}\cong s$ if $\hat{q}\ll1$. Furthermore, the standardization of **y** and small percent variance explained (PVE) imply $\hat{\sigma}\cong\sigma\cong1$. In our tests, *n* is sufficiently large that, under the null hypothesis *β* = 0, *t* is approximately standard normal with distribution function denoted by Φ. Rejecting the null hypothesis when the p-value is less than some critical value *α* is thus equivalent to rejecting when $\left| t \right|>-\Phi^{-1}\left( \frac{\alpha}{2} \right)\equiv z_{\alpha/2}\approx$5.0 for $\alpha=5\times{10}^{-7}.$

If *β* is actually nonzero, then $(\hat{\beta}-\beta)/\mathrm{se}(\hat{\beta})\cong(\hat{\beta}-\beta)\sqrt{s}$ is approximately distributed as standard normal and $t\sim N(\beta\sqrt{s},1)$. Therefore, the probability of rejection Pr$\left( \left| t \right|>z_{\alpha/2} \right)\approx$ $\Pr(N\left( \left| \beta\right|\sqrt{s},1 \right)>z_{\alpha/2})$ since $z_{\alpha/2}$ is so large. Therefore, the power to reject the null hypothesis is approximately $1- \Phi\left( z_{\alpha/2}-\left| \beta\right|\sqrt{2n\hat{q}} \right)$.
